# Long-read genomics reveal extensive nuclear-specific evolution and allele-specific expression in a dikaryotic fungus

**DOI:** 10.1101/2024.12.11.628074

**Authors:** Rita Tam, Mareike Möller, Runpeng Luo, Zhenyan Luo, Ashley Jones, Sambasivam Periyannan, John P. Rathjen, Benjamin Schwessinger

**Author notes:** Corresponding authors: John P. Rathjen and Benjamin Schwessinger.

## Abstract

Phased telomere to telomere (T2T) genome assemblies are revolutionising our understanding of long hidden genome biology “dark matter” such as centromeres, rDNA repeats, inter-haplotype variation, and allele specific expression (ASE). Yet insights into dikaryotic fungi that separate their haploid genomes into distinct nuclei is limited. Here we explore the impact of dikaryotism on the genome biology of a long-term asexual clone of the wheat pathogenic fungus *Puccinia striiformis* f. sp. *tritici*. We use Oxford Nanopore (ONT) duplex sequencing combined with Hi-C to generate a T2T nuclear-phased assembly with >99.999% consensus accuracy. We show that this fungus has large regional centromeres enriched in LTR retrotransposons, with a single centromeric dip in methylation that suggests one kinetochore attachment site per chromosomes. The centromeres of chromosomes pairs are most often highly diverse in sequence and kinetochore attachment sites are not always positionally conserved. Each nucleus carries a unique array of rDNAs with >200 copies that harbour nucleus-specific sequence variations. The inter-haplotype diversity between the two nuclear genomes is caused by large-scale structural variations linked to transposable elements. Nanopore long-read cDNA analysis across distinct infection conditions revealed pervasive allele specific expression for nearly 20% of all heterozygous gene pairs. Genes involved in plant infection were significantly enriched in ASE genes which appears to be mediated by elevated CpG gene body methylation of the lower expressed pair. This suggests that epigenetically regulated ASE is likely a previously overlooked mechanism facilitating plant infection. Overall, our study reveals how dikaryotism uniquely shapes key eukaryotic genome features.

## Introduction

Telomere-to-telomere (T2T) and haplotype-resolved genome assemblies have become the norm in eukaryotic genomics with advances in long-read sequencing technologies. Complete genome assemblies are fundamental for addressing key questions in eukaryotic genome biology that were previously intractable and hidden in the “dark matter” of genomes. Key breakthroughs have revolved around centromeres and the embedded kinetochore attachment sites pivotal for understanding karyotype diversity and evolution (Logsdon et al. 2024; Mastrorosa et al. 2024), as well as the notoriously repetitive ribosomal DNA (rDNA) arrays whose sequences have only been recently completed in humans (Nurk et al. 2022) and *Arabidopsis* (Fultz et al. 2023). Full resolution of haplotypes enables precise characterisation of inter-haplotype variations predominantly shaped by heterozygosity, structural rearrangements and transposable element movements (Ferguson et al. 2024; Gluck-Thaler et al. 2022; Hartmann 2022). Further, the complete picture of allelic information facilitates robust assessment of allele-specific expression (ASE) uncovering the underlying variations in *cis*-regulatory elements and epigenetic regulation, which can have important implications on phenotypic variability as well-established in mammals and plants (e.g. Shi et al. 2024; Cleary and Seoighe 2021; Tian et al. 2022; St. Pierre et al. 2022; Shao et al. 2019).

While there have been important novel insights into diploid and polyploid genome organisation, especially in plants (e.g. Belser et al. 2021; Sun et al. 2022; Hu et al. 2021), little is known about the fungi-specific dikaryotic state where two haploid genomes are contained in separate nuclei in the same cytoplasm and propagated in a coordinated manner during cell division (Anderson and Kohn 2014). Dikaryotism is highly successful with an estimated 400,000 taxa in the fungal subphylum Basidiomycota relying on it for significant time periods, for example during fruiting body formation in mushrooms and infection processes in rusts and smuts (James et al. 2020; Schmidt-Dannert 2016). Early partially-phased assemblies of rust fungi genomes indicated high levels of heterozygosity and presence/absence polymorphisms between the nuclear genomes, however they lacked the resolution to identify individual haplotypes (Schwessinger et al. 2020, 2018; Vasquez-Gross et al. 2020; Cuomo et al. 2017). The latest fully haplotype-phased and nuclear-assigned T2T genomes have provided the first insights into organisation of genes involved in mating behaviour and contributed to our understanding of reproductive mechanisms generating novel genetic diversity (Luo et al. 2024; Li et al. 2023a; Schwessinger et al. 2022). This includes the somatic exchanges of nuclei between asexual rust lineages that are adapted to the infection of cereals (Henningsen et al. 2024; Sperschneider et al. 2023), and sexual recombination in permissive hosts (Wang et al. 2022; Rodriguez-Algaba et al. 2022; Du et al. 2023). Such genetic reshuffling produces novel allele combinations in so-called avirulence (*Avr*) genes that encode secreted effector proteins essential for host infection. Rust effectors are under strong selection pressure to diversify into non-recognised virulence alleles because they can be recognised by cognate immune receptors encoded by wheat resistance genes (Salcedo et al. 2017; Ortiz et al. 2022; Chen et al. 2017). This is necessary to escape the plant immune system and to confer the ability to infect new host varieties.

Key questions remain around the impact of dikaryotic genome organisation on the evolution of centromeres, rDNA repeats, detailed inter-haplotype variations and ASE. Here we use a long-term asexual clone of the wheat stripe rust fungus *Puccina striiformis* f. sp. *tritici* (*Pst*) dating back at least 80 years to address these questions in a globally important wheat pathogen (Schwessinger et al. 2018; Wellings 2007; Thach et al. 2016). We leveraged high-accuracy Oxford Nanopore Technology (ONT) duplex long-reads to generate the first ONT-based T2T, fully nuclear-phased genome assembly for a dikaryotic fungus. We combined this with comprehensive long-read ONT cDNA datasets sampled during dormancy and pathogenesis for high-quality gene annotations and differential expression analysis at gene and allele-specific level. Our study sheds new light onto the genome biology and adaptive evolutionary potential of rust fungi, with important implications for managing their agricultural impacts.

## Results

### A T2T nuclear-phased genome assembly of *Puccinia striiformis* f. sp. *tritici* 104E based on high-quality ONT long-read sequencing

We set out to generate the first T2T genome assembly for the dikaryotic fungus *Puccinia striiformis* f. sp. *tritici* using only ONT long-read sequencing while making use of high-accuracy duplex data. We sequenced DNA extracted from dikaryotic urediniospores of a representative isolate of the Australian founder pathotype 104E137A- (abbreviated as *Pst*104E), which belongs to the long-term asexual *Pst*S0 lineage (Schwessinger et al. 2018, 2020). We used Verkko (Rautiainen et al. 2023) to assemble duplex reads (>10 kbp and >Q30; ∼32x sequencing depth per haplotype) combined with ultralong (UL) simplex reads (>40 kbp and >Q10; ∼117x sequencing depth per haplotype) followed by Hi-C phasing. The raw assembly consisted of 355 contigs and one scaffold totalling 168 Mbp, with 39 contigs covering 90% of the total assembly length, indicating high contiguity (Supplemental Table S1). We used Hi-C linkage information to join fragmented contigs into three additional chromosomes (chr), resulting in 36 chromosomal scaffolds corresponding to the 18 homologous pairs, which we sorted by average length (chr1 to chr18) (Fig. 1A; Supplemental Fig. S1). A Hi-C contact heatmap clearly grouped the two sets of 18 chromosomes separately, each representing one nuclear complement, as expected for dikaryotic fungi (Fig. 1B). We also curated the assembly using manual procedures such as gap filling and telomere recovery (Supplementary Notes). This resulted in 35 T2T chromosome assemblies, with only chr16A missing its q-arm telomere. The 71 assembled telomeres had a mean length of 259 bp, corresponding to ∼43 repeats per telomere, in line with other basidiomycete fungi (Ramírez Nasto et al. 2011; Schwessinger et al. 2020; Sperschneider et al. 2021). Self-alignment dotplots and the assembly graph revealed sources of the five remaining gaps: a ∼650 kbp retrotransposon-dominated region on chr6A residing near the previously identified mating type *PR* locus (Luo et al. 2024) (Supplemental Fig. S2); two unresolved regions on chr8A as hinted by graph bubbles (Supplemental Fig. S3); and two rDNA arrays on chr13A and 13B.

**Figure 1.**
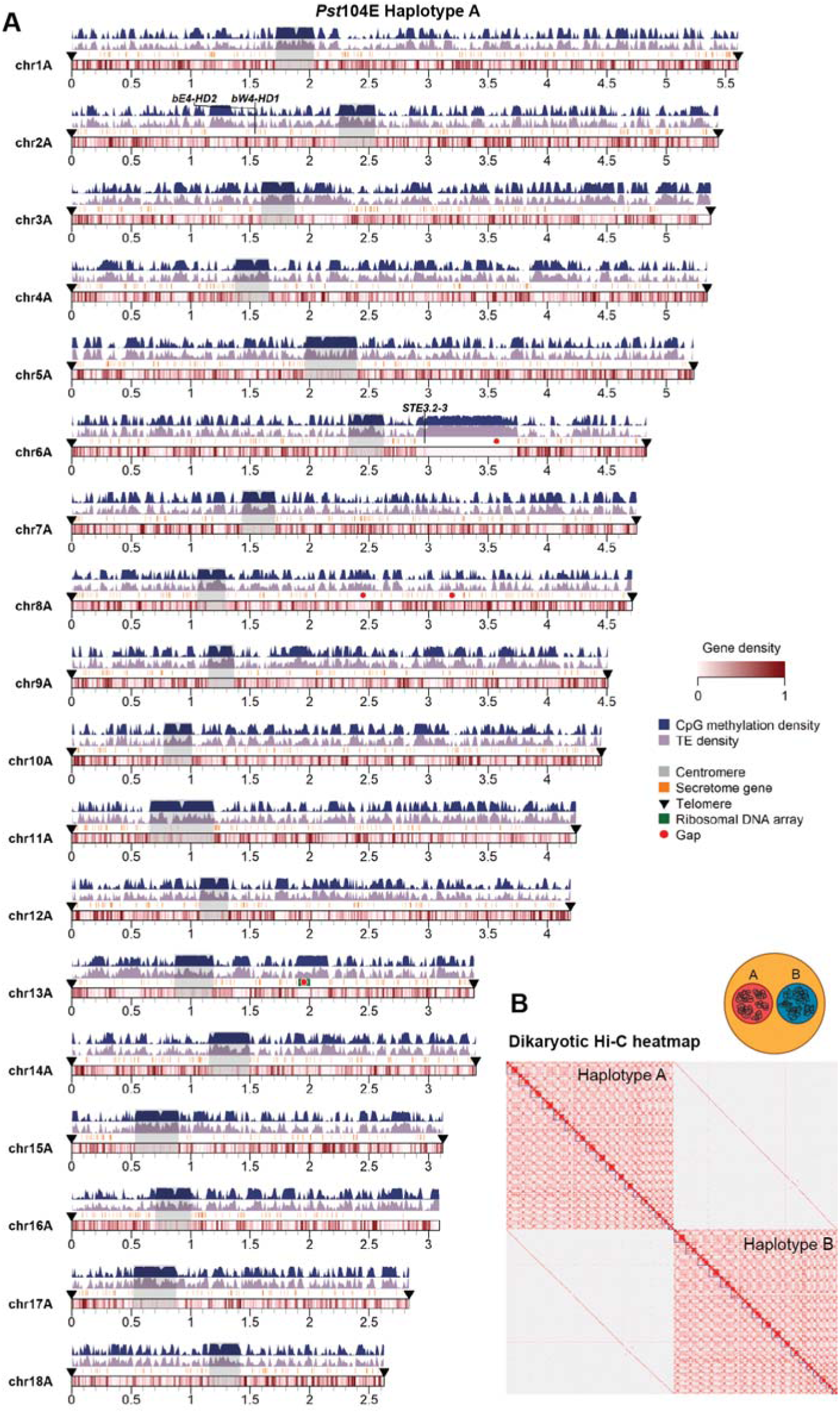
A nuclear-phased, chromosome-scale genome assembly of the dikaryotic fungus *Puccinia striiformis* f. sp. *tritici* isolate *Pst*104E. **(A)** Karyoplot of the 18 chromosomes of haplotype A, showing density of CpG methylation and transposable elements (TEs) as peaks, and gene density as heatmaps within chromosome ideograms (10 kbp sliding windows). Locations of centromere, telomeres, secretome genes, ribosomal DNA array and assembly gaps are annotated as per legend. **(B)** Hi-C contact heatmap of the full dikaryotic genome assembly consisting of 36 chromosomes. The nuclear haplotypes A and B display a clear signal of spatial separation.

We evaluated the quality of the final curated assembly in terms of completeness of core genes, contiguity, and correctness. The full dikaryotic assembly yielded 92.6% of complete BUSCOs of the Basidiomycota lineage (Simão et al. 2015), equivalent to recently published PacBio HiFi genome assemblies of other dikaryotic rust fungi (Wang et al. 2024; Li et al. 2023a; Sperschneider et al. 2023). We used the long terminal repeat (LTR) assembly index (LAI) (Ou et al. 2018) to assess contiguity at repetitive LTR retrotransposons. Haplotype A and B assemblies each received LAI scores of 27.2 and 24.8, classifying them to the highest rank based on well-assembled repeats. Merqury (Rhie et al. 2020) k-mer-based quality value was 57.4, corresponding to >99.999% consensus accuracy, in line with recent PacBio HiFi assemblies (Wang et al. 2024; Li et al. 2023a; Sperschneider et al. 2023). CRAQ (Li et al. 2023b) was used to detect clipped alignments indicative of regional and structural errors that could be computed into R/S- Assembly Quality Indices (AQIs). The assembly achieved R/S-AQI of 94.7 and 97.4, both meeting reference quality. Rare residual errors were reflected by the low density of SNPs detected as homozygous (1.1/Mbp, 161 total) and heterozygous (6.8/Mbp, 1038 total).

Analysis of the mapping read depth supported full haplotype phasing with a single peak corresponding to the expect haploid 1x depth (Supplemental Fig. S4). To detect the presence of putative phase switch errors, we quantified Hi-C paired alignments (MAPQ ≥ 20) within- and between-haplotypes over 20 kbp genomic windows. About 99.2% of the Hi-C mappings were between chromosomes contained within one nucleus (within-haplotype links), with less than 0.8% linking chromosomes contained in separate nuclei (cross-haplotype links) (Table 2). This suggests that our assembly had only very few residual phase switch errors. Visualising Hi-C links on a circos plot showed evenly distributed, low-frequency cross-haplotype Hi-C links across genomic windows which could be attributed to noisy Hi-C read alignments (Supplemental Fig. S5). An exception could be seen in a chr11B window showing abnormally high number of cross-haplotype links, potentially identifying a minor residual phase switch. These results demonstrate a highly complete and accurate nuclear-phased genome assembly of *Pst*104E.

### ONT cDNA sequencing enables high-quality evidence-guided gene annotations

We aimed to improve on current fungal gene annotations by incorporating extensive long-read cDNA sequencing datasets (Pardo-Palacios et al. 2024). We generated a detailed time-course of ONT direct cDNA datasets for *Pst*104E gene annotation and differential expression analysis. The transcripts were sampled from six conditions with four replicates each (Supplemental Table S2). We sampled ungerminated (UG) urediniospores as the dormancy control, and infected wheat leaf tissues at 4-, 6-, 8-, 10- and 12-days post infection (dpi). Principal component analysis (PCA) demonstrated clear clustering of the technical replicates of each sample and revealed separation of samples representing UG, early- (4 and 6 dpi) and late-infection (8, 10 and 12 dpi) conditions (Supplemental Fig. S6). We complemented gene annotations with multiple publicly available Illumina RNA-seq datasets (Dobon et al. 2016; Schwessinger et al. 2018; Zhao et al. 2021) (Supplemental Table S3). As the long- and short-read datasets were collected from different biological samples, we independently processed them using different bioinformatic tools (see Methods) and supplied them to the fungi-tailored gene annotation pipeline funannotate (Jonathan and Jason 2023). In total, we annotated 15,142 genes on haplotype A and 14,938 protein-coding genes on haplotype B. Our annotated gene space improved the complete BUSCO score to 94.8% highlighting the quality of our annotation approach. Functional annotation identified 2,249 and 2,288 genes, for haplotypes A and B respectively that encode secreted proteins without predicted transmembrane domain.

We identified potential candidate *Avr* effector genes in *Pst*104E via differential gene expression analysis using our ONT cDNA dataset. We searched for secretome genes that are upregulated early during wheat infection (4 and 6 dpi) relative to UG, as their functions likely correlate with pathogenesis. We found a total of 1,105 secretome genes that were upregulated early during infection. These were shortlisted to hemizygous genes for reduced functional redundancy, resulting in 89 high-priority candidates for future functional validation (Supplemental Table S4).

### Transposable element annotations

Transposable elements (TEs) are major components of genomes including many basidiomycete fungi (Castanera et al. 2017). We identified TEs for each haplotype and classified them to the superfamily level using the REPET pipeline (Quesneville et al. 2005; Flutre et al. 2011). Both haploid genomes had similar TE content and composition, covering ∼44.51% of the genome space (Table 1). This was consistent with findings in other published *Pst* genomes (Schwessinger et al. 2018; Zheng et al. 2013; Schwessinger et al.) . Class I (retrotransposons) and Class II (DNA transposons) accounted for 15% and 18.7% of the genome respectively (Supplemental Table S5). LTRs comprised the most abundant retrotransposons, predominated by the Ty3/Gypsy. Terminal inverted repeats (TIRs) represented the majority of DNA transposons.

**Table 1.**
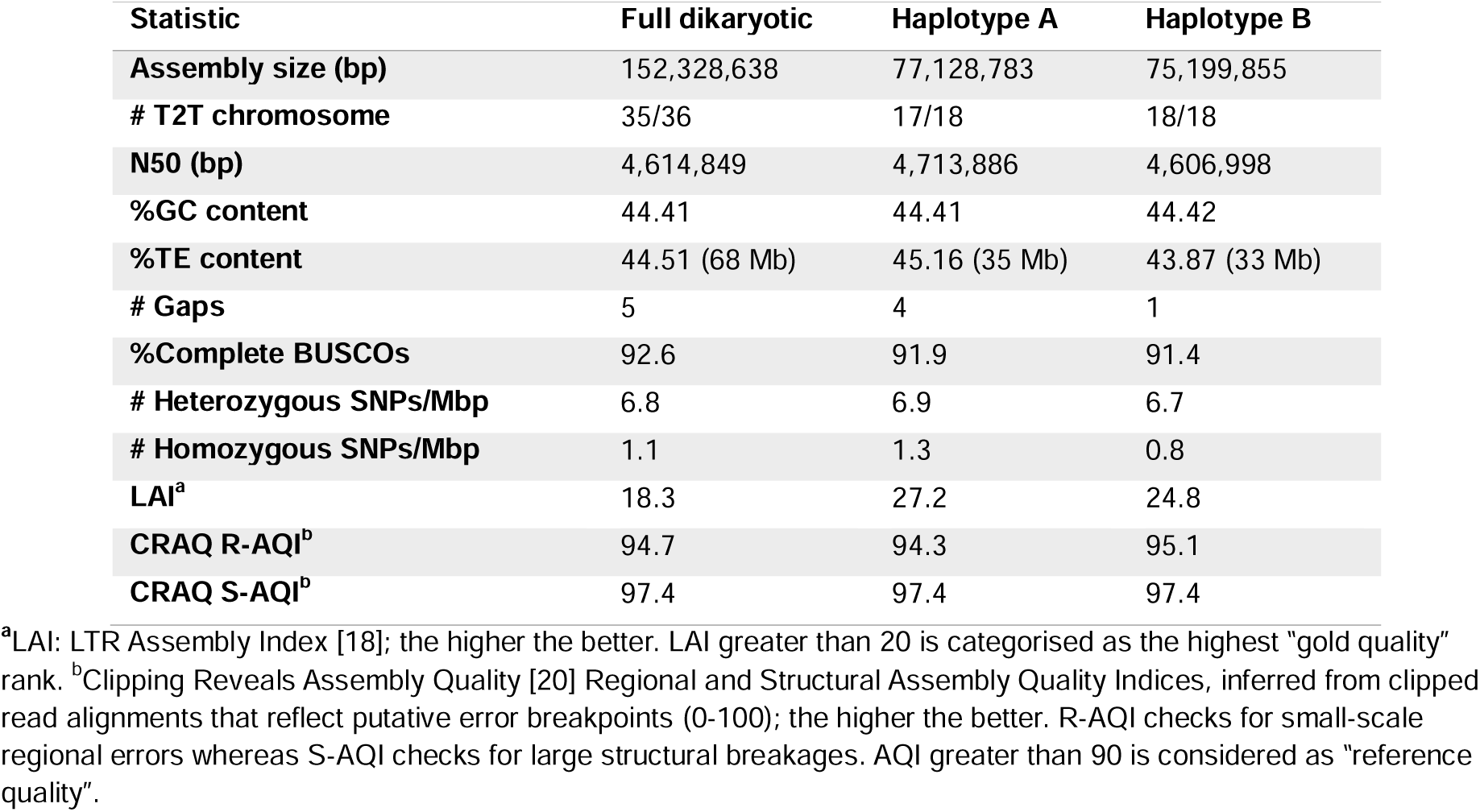
Summary of assembly statistics and quality metrics of the ONT duplex genome assembly of *Pst*104E.

**Table 2.** Hi-C contact statistics confirming phasing correctness of the *Pst*104E genome assembly.

| Types of Hi-C contacts (MAPQ ≥ 20) | Counts |
| --- | --- |
| **Total Hi-C links** | **1,126,566** |
| **Within-haplotype links** | **1,117,107 (99.2%)** |
| ***cis*-chromosome** |  |
| Haplotype A | 479,583 (42.6%) |
| Haplotype B | 467,937 (41.5%) |
| ***trans*-chromosome** |  |
| Haplotype A | 85,667 (7.6%) |
| Haplotype B | 83,920 (7.5%) |
| **Cross-haplotype links** | **9,459 (0.8%)** |

### *Pst* centromeres are highly diverse and enriched with LTR retrotransposons

We set out to identify *Pst*104E’s centromeric regions and investigate their sequence composition. Basidiomycete centromeres are often characterised by hypermethylated TE-rich and gene-poor regions (Guin et al. 2020). To define *Pst* centromeres, we first estimated their positions from Hi-C contact heatmaps based on the distinctive bowtie-like structures classic for centromere-to-centromere interactions (Fig. 1B) (Varoquaux et al. 2015; Muller et al. 2019). We investigated the sequence composition and methylation status of these genomic regions. The bowtie-like interaction sites were enriched for TEs and overlapped with gene-sparse regions spanning hundreds of kilobases(Fig. 1A, 2A). We next determined the CpG methylation status using our ONT long-read sequencing dataset. We found substantially more methylated CpG within all 36 inferred centromeric regions, *Cen1A* to *Cen18B* (94.9% to 98.3%), than in non-centromeric regions (32.9% to 45.3%) (Fig. 2B).

**Figure 2.**
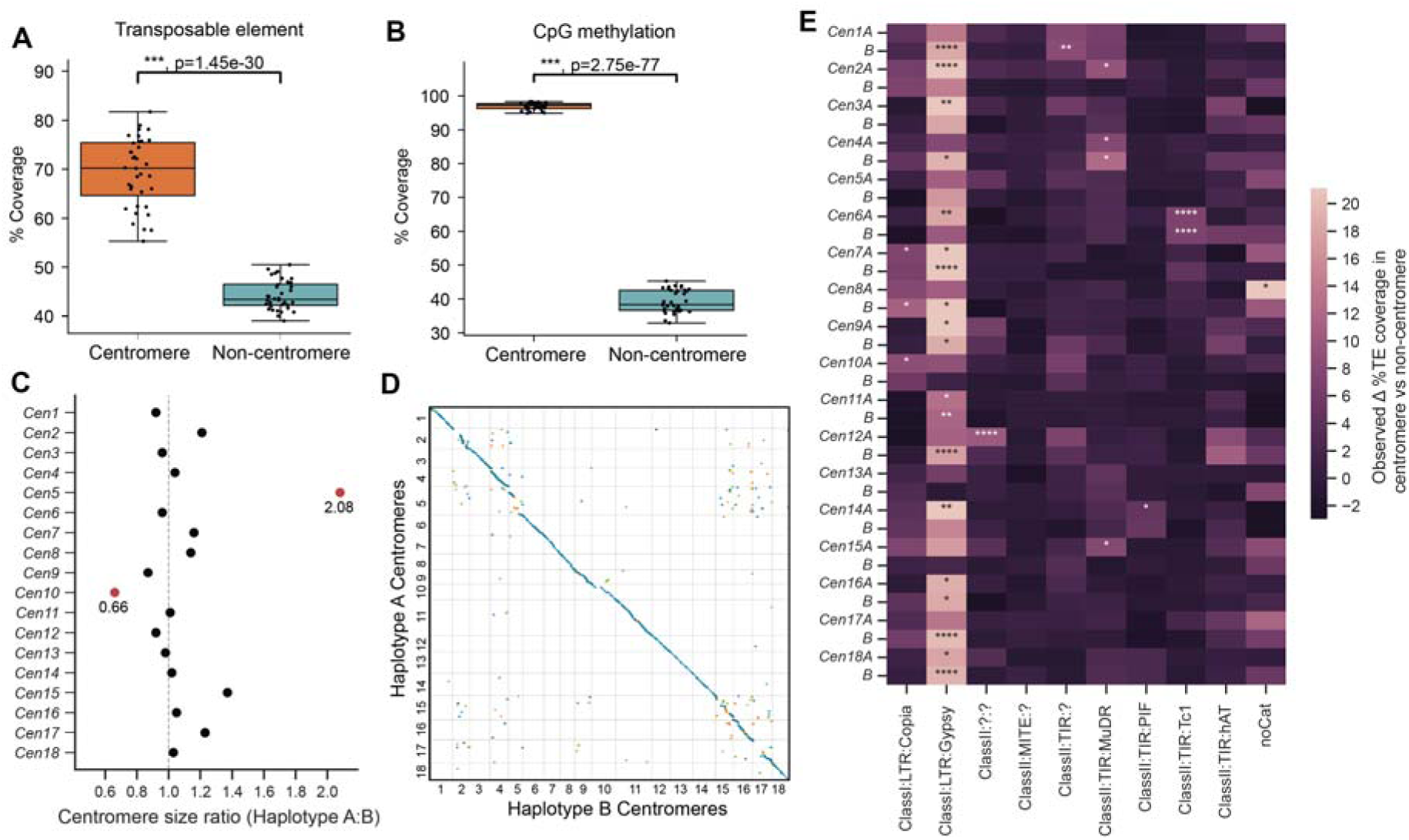
*Pst*104E centromeres are highly diverse with haplotype-specific sequences and are enriched in retrotransposons. **(A)** shows the percentage sequence coverage distribution of centromere and non-centromere regions of *Pst104E*. Each dot represents one of 36 *Pst104E* chromosomes. Student’s *t-*test (***, p < 0.001). **(B)** shows the percentage coverage of methylated CpG sites in centromere and non-centromere regions of *Pst104E*. Each dot represents one of 36 *Pst104E* chromosomes. Student’s *t-*test (***, p < 0.001). **(C)** shows the size ratio of haplotype A centromeres compared to haplotype B centromeres. Outlier ratios (1.5-times the interquartile range) are highlighted in red. **(D)** NUCmer pairwise alignment dotplot between centromeres of haplotypes A and B. Only alignment blocks longer than 100 bp with minimum 90% sequence identity are shown. Each colour denotes an alignment type: blue, unique forward alignments; green, unique reverse alignments; orange, repetitive alignments. **(E)** shows the enrichment analysis of different transposable elements (TE) within centromeres when compared with non-centromere regions. TE superfamily enrichment at centromeres was statistically tested with permutation tests for each chromosome. Only abundant TE superfamilies with >1% total genome coverage are shown. The colour scalebar denotes the test statistic, which is differences in % TE coverage within and outside centromeres. P-value represents the proportion of expected values as or more extreme than observed. FDR < 5% applied to correct for multiple testing (*, p<0.05; **, <0.01; ****, <0.0001).

*Pst*104E centromere sizes varied approximately 2.5-fold between 210 and 538 kbp (mean 304 kbp; Supplemental Table S6), categorising them as “large regional” centromeres which are known to support multiple spindle microtubule attachment sites (Yadav et al. 2018a). Most *Pst*104E homologous chromosomes share similar centromere lengths between haplotypes with length differences under 1.4-fold, while *Cen5A* stood out with double the length of *Cen5B* (Fig. 2C).

Pairwise alignments of centromeric regions revealed varying levels of macro-collinearity between haplotypes, ranging from almost complete (e.g., *Cen3, 12, 13* and *18*) to negligible synteny (e.g., *Cen2, 5, 15* and *17*) (Fig. 2D). Given the high TE density, such a diverse range in centromeric sequence conservation prompted us to examine their TE composition in search of elements possibly linked to centromere function.

We conducted permutation tests to determine the enrichment of abundant TE superfamilies within each *Pst* centromere compared to non-centromeric regions (Fig. 2E; Supplemental Table S7). Most centromeres (25 out of 36) were found to be significantly enriched for one to two TE superfamilies that belonged to retro- and/or DNA transposons (Fig. 2E). Of these, 19 centromeres were significantly enriched for Ty3/Gypsy LTRs, occasionally co-colonised by TIRs. Given the balanced representation of retro- and DNA transposons throughout the genome, this finding suggests that retrotransposons might have a more prominent role in *Pst* centromere formation than DNA transposons.

### Centromeres contain a single putative kinetochore attachment site

It is unclear if rust fungi have one or multiple kinetochore attachment sites given they have “large regional” centromeres (Figure 2) (Yadav et al. 2018a; Sperschneider et al. 2021). Kinetochores typically assemble at a hypomethylated stretch of DNA embedded within the centromere termed the “centromere dip region” (CDR) which is marked by the centromere-specific histone variant CENP-A (Logsdon et al. 2024; Sundararajan and Straight 2022; Akiyoshi 2019). We examined the CpG methylation pattern along the *Pst*104E centromeres to locate CDRs as a qualitative proxy for potential kinetochore sites. Throughout all centromeres, we consistently observed a single methylation depletion “valley” spanning 24.8 Kbp on average, which is characteristic of CDR (Fig. 3A; Supplemental Fig. S7; Supplemental Table S6). The CDR lengths were similar between haplotypes (Fig. 3B). Interestingly, the positioning of CDRs were generally conserved (Fig. 3C, 2D). The two notable exceptions were *Cen3* and *Cen13*, whose CDR haplotypes were placed further than 20% of their centromere lengths apart (Fig. 3C).

**Figure 3.**
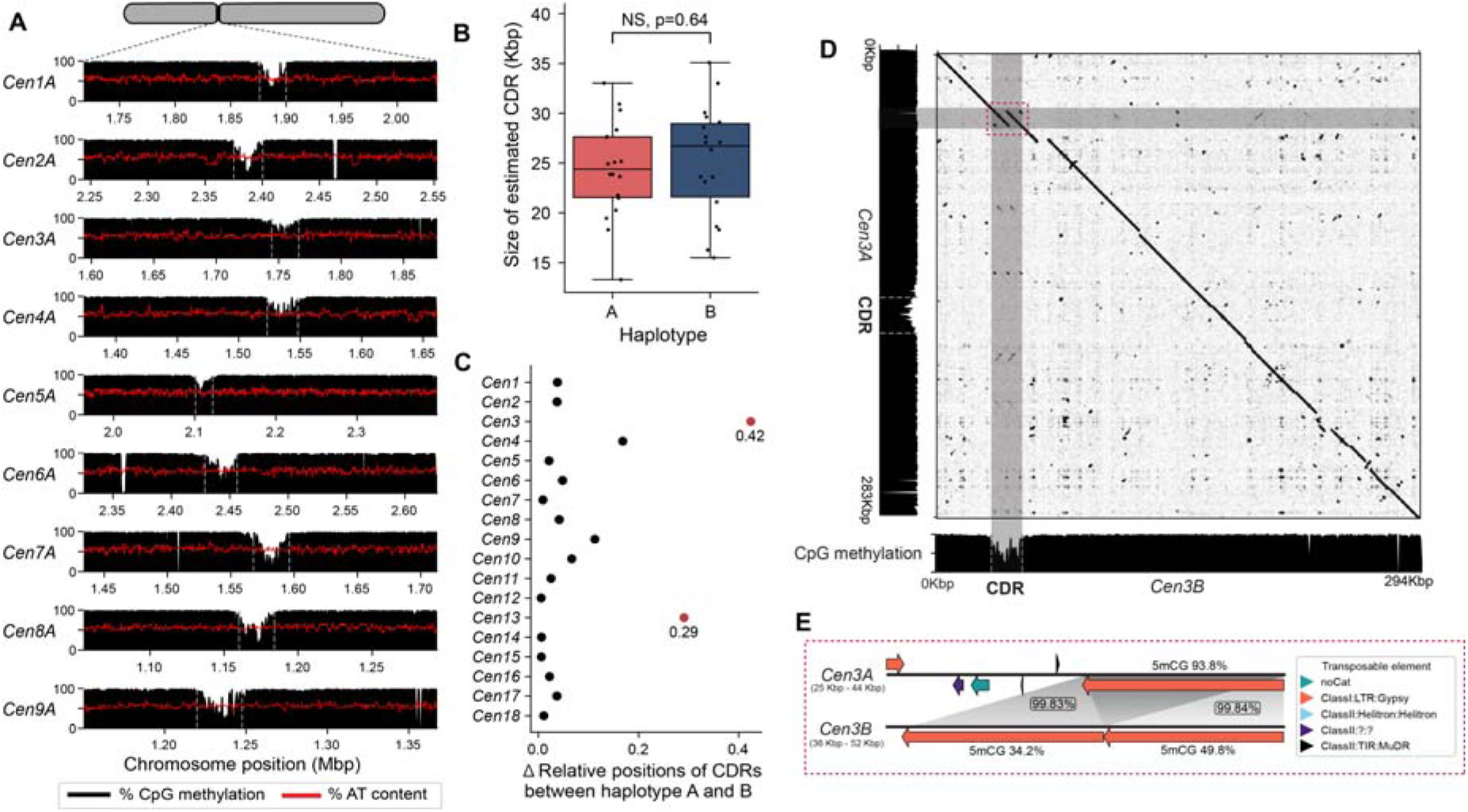
Analysis of centromere dip regions (CDRs) of Pst104E centromeres suggest a single kinetochore attachment sites per chromosome. **(A)** shows the CpG methylation profiles (black histograms) and percentage AT content (red line) of *Pst*104E centromeres 1A to 9A as examples. A methylation depletion valley with the mean size of ∼24.8 Kbp consistently appeared throughout all centromeres, indicating CDR signals (marked by grey dotted line). **(B)** shows the estimated sizes of the CDRs for each chromosome in haplotype A and B. Student’s *t-*test (NS, p > 0.05). **(C)** shows the differences in the relative positions of the CDRs between haplotypes for each chromosome pair. Differences exceeding 0.2 are highlighted in red. **(D)** shows the sequence alignment dotplot between *Cen3A* and *3B* and their respective CpG methylation profiles (black histograms). The dotted red box highlights the sequence divergence between *Cen3A* and *3B* corresponding to the *Cen3B* CDR only. Visualised in gepard [28]. **(E)** shows detailed synteny analysis of the red box highlighted region of (D). The long orange arrows indicated copies of hapB-B-G1437-Map9 belonging to the Ty3/Gypsy LTR retrotransposon superfamily. Grey shading indicates homologous sequences with the percentage identity shown in boxes. The numbers above and below the long orange arrows indicate the respective percentage coverage of methylated CpG sites.

We next investigated whether the CDR signals were linked to increased AT content as reported for other fungi (Narayanan et al. 2024; Yadav et al. 2018a; Sankaranarayanan et al. 2020). No differences in AT content were detected when we compared CDRs with other centromeric or non- centromeric regions (Supplemental Fig. S8). The only exception was *Cen3B* CDR, which has an increased AT content of 59.8% when compared to the rest of the centromere and the overall genome average of 55.6%. Close investigation revealed two nearly identical AT-rich copies of a Ty3/Gypsy retrotransposon family (hapB-B-G1437-Map9), together covering 91% of *Cen3B* CDR (Fig. 3D). Haplotype sequence alignment revealed that the corresponding region on *Cen3A* contained only one copy of this TE family which is not involved in CDR formation. Consistently, the single copy TE insertion of hapB-B-G1437-Map9 on *Cen3A* was highly methylated on CpG sites (93.8%) while the two TE copies on the CDR of *Cen3B* were lowly methylated (34.7% and 49.8%). This is despite over 99.5% sequence identity shared by the three TE copies. This highlights the fact that centromere and kinetochore attachment site formation is not solely driven by primary DNA sequence composition.

### T2T haplotype-resolved genome assembly reveals nucleus-specific variations in the rDNA arrays

We investigated the rDNA composition in *Pst*104E to better understand its dynamics in the context of a dikaryotic genome with physical separation ofthe haploid genomes. *Pst*104E has a single rDNA cluster per nuclear genome on the q-arm of chr13 (Fig. 1A). Both haplotypes were incompletely assembled with a gap surrounding the rDNA cluster indicating an underrepresentation of this complex locus (Supplemental Fig. 3). The assembled rDNA copies are oriented such that the transcription direction points away from the centromere and towards the telomeric region (Fig. 4A). To define a canonical full-length rDNA repeat, we aligned reference ITS and rRNA genes of several *Puccinia spp.* (see Methods) supplemented with our rRNA long reads (Supplemental Fig. S9). This allowed us to reconstruct a canonical *Pst* rDNA repeat including the 45S transcription unit, which consists of 18S, 5.8S and 25S rRNA genes, separated by two internal transcribed spacers (ITS1 and ITS2) (Fig. 4B). This is followed by two intergenic spacers (IGS1 and IGS2), separated by a 5S rRNA gene that is barely transcribed. We defined 18S and IGS2 including the 5’ external transcribed spacer regions as the start and end of an rDNA repeat unit for the subsequent analysis (Fig. 4B).

**Figure 4.**
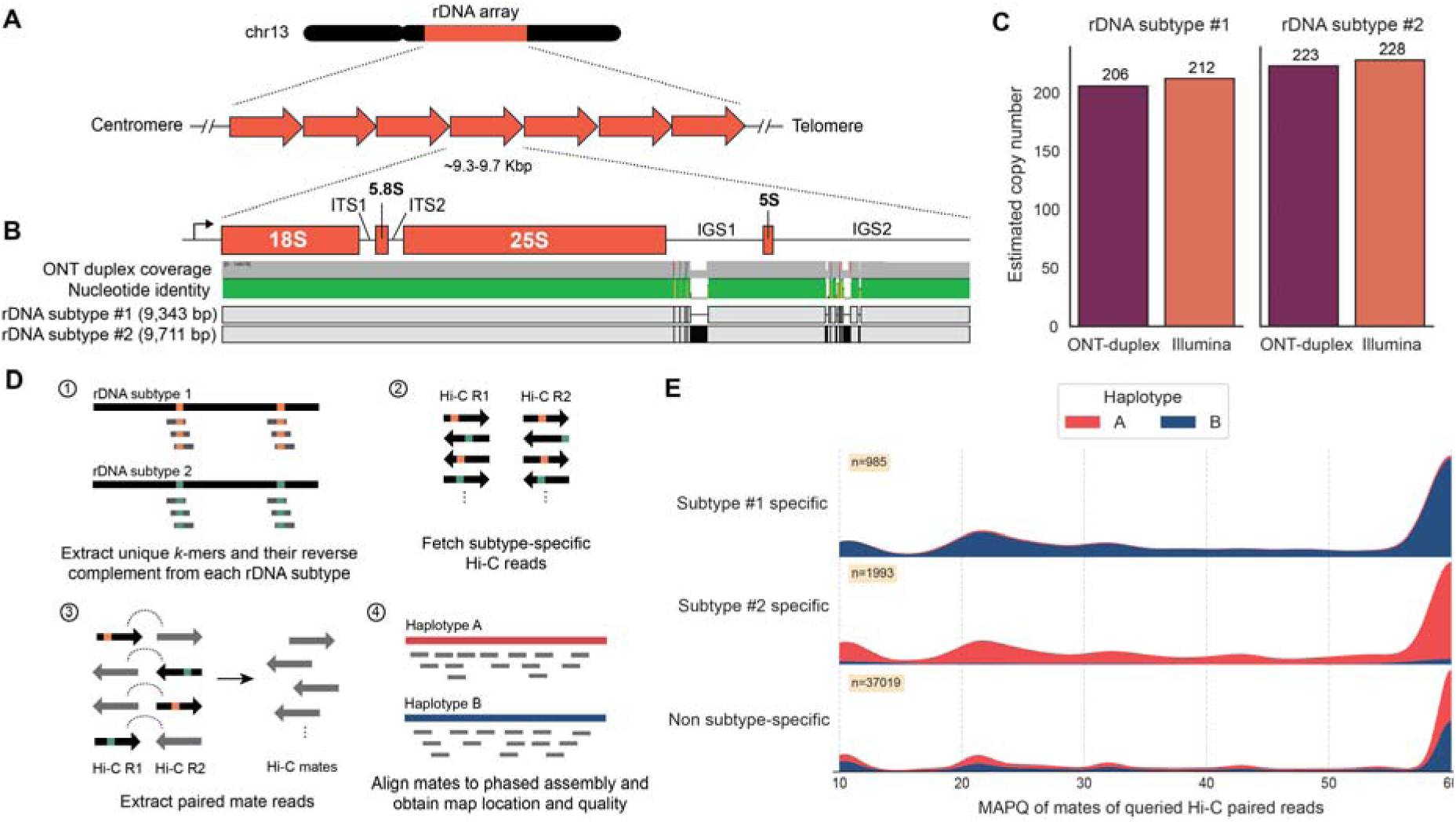
The two dikaryotic nuclear haplotypes of *Pst104E* harbour their own unique rDNA subtypes. **(A)** shows a schematic view of the rDNA tandem repeat array located on chromosome 13. **(B)** shows a diagram of a single canonical rDNA unit of *Pst*104E containing the transcription start site in the 5’ external transcribed spacer (5’ ETS; not shown), the catalytic rRNA genes (18S, 5.8S, 25S and 5S), two internal transcribed spacers (ITS1 and ITS2) and two intergenic spacers (IGS1 and IGS2). The alignment indicates two major rDNA subtypes containing SNPs and structural differences in IGS1 and 2. **(C)** shows the estimate coverage of rDNA subtype #1 and #2 based on ONT-duplex and Illumina raw read datasets. **(D)** describes the detailed workflow of our rDNA Hi-C analysis designed to determine the physical location of the rDNA subtypes in each nuclear haplotype. **(E)** shows the mapping quality (MAPQ) distributions of Hi-C reads whose pair contained rDNA subtype-specific or non-subtype-specific 31-mers, categorised by the haplotype they mapped against.

We examined the intra-isolate rDNA sequence variations by mapping duplex reads against the canonical rDNA unit defined above. This revealed two major rDNA subtypes supported by substantial read coverage at near equal frequencies. The two subtypes (#1 and #2) are 9,343 bp and 9,711 bp in length, with polymorphisms contained in the repeats nested within IGS1 and IGS2 (96.1% sequence identity; Supplemental Fig. S10). We also identified twelve rarer variants of the two subtypes (#1.1-1.9 and #2.1-2.3) based on low-frequency SNP analysis (Supplemental Fig. 11 and Supplemental Table S8). Intriguingly, most of these SNPs occurred in the 18S gene.We estimated the copy number of each rDNA subtype by their ONT and Illumina sequencing depths (Schwessinger et al. 2018; Lofgren et al. 2019; Sharma et al. 2022). Both supported similar estimates, with 206 to 212 copies for rDNA subtype #1 and 223 to 228 copies for subtype #2, totalling approximately 434 copies (Fig. 4C). The subtype variants #1.1-1.9 and #2.1-2.3 collectively accounted for about 23% of the total rDNA copy number, suggesting incomplete rDNA homogenisation (Supplemental Table S8).

Given the long clonal history of *Pst*104E (Schwessinger et al. 2020), the individual haplotypes are expected to have been stably inherited in separate nuclei without karyogamy or meiotic crossovers. We therefore hypothesised that the sequence variations in the two dominant rDNA subtypes might be nucleus-specific. To test this, we took advantage of Hi-C read pairs that contained rDNA subtype-specific k-mers and asked where the alternate read mate pair mapped (Fig. 4D). About 91.9% of mates associated with rDNA subtype #1 mapped to haplotype B, whereas 93.3% of those associated with subtype #2 mapped to haplotype A (Fig. 4E). The control procedure in which we mapped subtype-nonspecific rDNA Hi-C read pairs displayed a near-equal proportion of Hi-C mates mapped against each haplotype. Together, this implies that each rDNA subtype was associated with a different haplotype and that each nuclear haplotype contains its own major rDNA subtype array.

### Large scale structural variations driven by transposable elements shape the inter-haplotype diversity of *Pst104E*

The phased *Pst*104E assembly allowed us to assess its inter-haplotype structural variations (SV). The haplotypes were about 80% syntenic (Fig. 5A), with 60.2 Mbp of haplotype A (78.0%) and 60.1 Mbp of haplotype B (79.6%) identified as highly continuous syntenic blocks (Fig. 5B). We identified a total of 1,734 duplications, 1,024 indels (≥50 bp), 408 translocations and 19 inversions, cumulatively occupying about 11% and 9% of the total lengths of haplotypes A and B, respectively (Fig. 5B; Supplemental Table S9). About 10% of each haplotype’s sequence was not alignable and is therefore hemizygous. Syntenic regions had the largest average length with most ranging between 10 and 100 kb. The different SV types had various length distributions mostly centred around 1-10 kb (Fig. 5B). We used permutations to test if SVs and their 2 Kbp flanking regions were enriched for specific genomic features. This revealed a significant enrichment of TEs across all analysed SV types, especially duplications, possibly due to the replicative mechanism of retrotransposons (Fig. 5C; Supplemental Table S10). In contrast, protein-coding genes were depleted at SVs.

**Figure 5.**
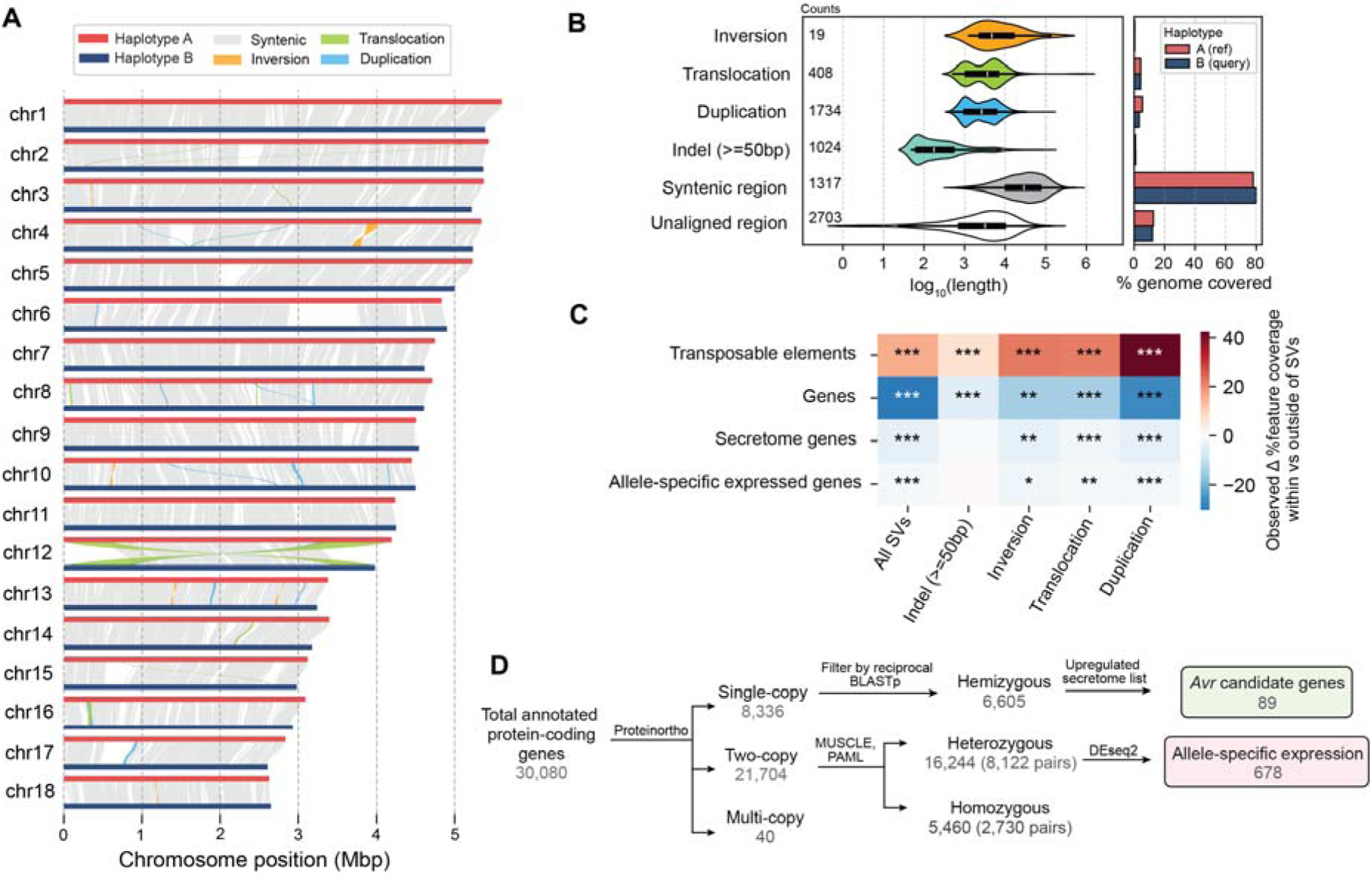
*Pst104E’s* inter-haplotype variations is shaped by large scale structural variation and transposable elements. **(A)** shows the synteny and structural rearrangements between chromosome pairs of the two nuclear haplotypes, with A as reference and B as query. **(B)** shows the length distribution (left) of different structural variation (SV) types, including syntenic and unaligned regions. The counts of each category are provided. The bar chart (right) represents the total genome length covered by each SV type and region. **(C)** shows the enrichment (red) and depletion (blue) of different genomic features (TEs and genes) within SVs and in close proximity (+/- 2 kbp). The colour scale denotes the difference in total percentage coverage of a genomic feature. P-values indicates the proportion of expected genomic feature coverage less than, at, or more than the observed value (*, p<0.05; **, <0.01; ***, <0.001; FDR-corrected). **(D)** shows the workflow summarising the classification of hemizygous, heterozygous and homozygous protein-coding genes, and their further categorization. Hemizygous genes were intersected with secretome genes upregulated during infection timepoints to shortlist *Avr* candidates. Heterozygous biallelic genes were used to analyse allele-specific expression.

Next, we assessed the impact of inter-haplotype variation on protein-coding genes by analysing sequence conservation and synteny (Fig. 5D). Using Proteinortho (Lechner et al. 2011) and reciprocal BLAST searches, we robustly identified 3,391 and 3,214 hemizygous (single copy) genes unique to either haplotype A or B, respectively. This is consistent with about 20% of each haplotype’s sequence being hemizygous. A total of 21,744 genes share at least one homolog in the alternative haplotype. We analysed one-to-one gene pairs and performed codon-aware alignment to compute their divergence values at synonymous and non-synonymous sites (Supplemental Table S11; see Methods). This process identified 2,730 homozygous pairs. The remaining 8,122 displayed divergence greater than zero, thereby are defined as heterozygous biallelic pairs.

### Allele-specific expression is correlated with gene body methylation profiles and enriched for genes predicted to be implicated in plant infection

We next investigated if any of the heterozygous allele pairs displayed allele-specific expression (ASE) in resting spores (UG) or during the wheat infection process at five different time points (4, 6, 8, 10, 12 dpi).

We first tested if there is an overall expression bias at the individual nuclear haplotype level. Both haplotypes displayed balanced gene expression without evidence of nuclear dominance (Supplemental Fig. S12); however, an expression bias for haplotype A was detected when considering only the heterozygous biallelic genes (Fig. 6B).

**Figure 6.**
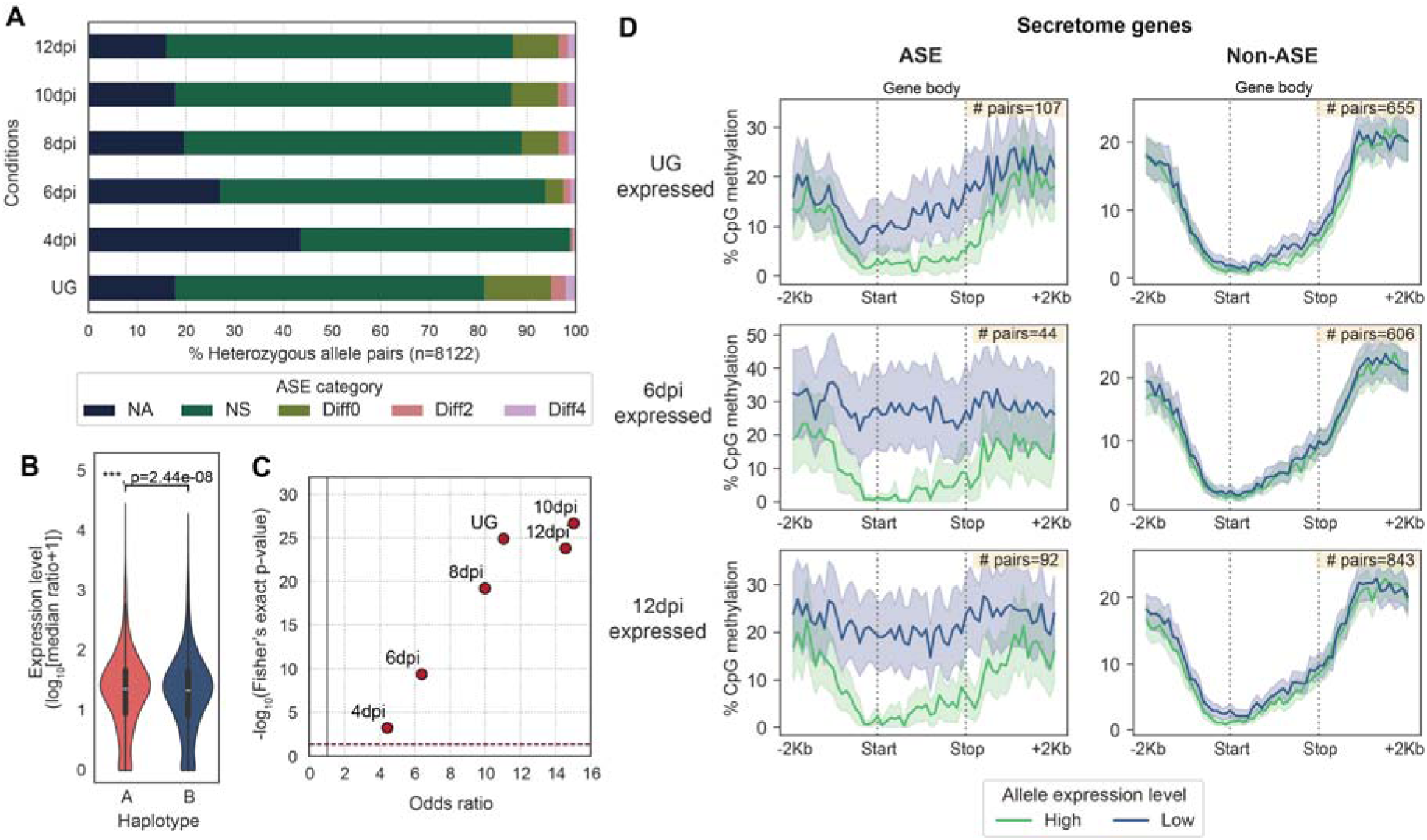
Allele-specific expression (ASE) is prevalent for secretome genes and correlated with gene methylation patterns. **(A)** shows the five ASE categories detected across six different transcript sampling conditions. Different colors indicate ASE categories. Allele pairs displaying log2 fold changes greater than two (Diff2 and Diff4) in at least one condition were defined as ASE in subsequent analysis. **(B)** shows the expression level of alleles belonging to the two nuclear haplotype A and B. Mann-Whitney U test (***, p < 0.001). **(C)** shows the odds ratio and log_10_ of the Fisher’s exact text p-value for ASE genes comparing secretome genes with evolutionarily conserved BUSCOs. Each red dot represents a transcript sampling conditions as labelled. The dotted red and grey line highlights cut-offs matching the null-hypothesis of no difference between the two gene groups. **(D)** shows the distribution of CpG methylation (sampled at UG) across secretome gene bodies (here defined as start to stop codon) including +/- 2 Kbp flanking regions. The mean percentage coverage CpG methylation is shown as solid line; 95% bootstrapping confidence interval as shaded area. The yellow inset indicates the number of allele pairs included.

We compared the transcript abundance for each heterozygous allele pair to determine its ASE across all six tested conditions (Fig. 6A, Supplemental Table S12). We classified the ASE status using the following criteria (Shi et al. 2024): (1) no observed expression or no unambiguous transcript mapping at both alleles (NA); (2) no significant difference between alleles with false discovery rate (FDR) adjusted p-value > 0.05 (NS); and (3) significant difference between alleles with adjusted p-value < 0.05, indicating differential ASE. The differential ASE pairs were further classified based on log_2_ fold change (LFC): weak ASE, |LFC|<2 (Diff0); moderate ASE, 2≤|LFC|<4 (Diff2); and strong ASE, |LFC|>4 (Diff4). While the majority of heterozygous gene pairs showed no evidence of ASE, a significant proportion exhibited ASE in each condition. The UG stage had most ASE pairs (18.7%), followed by 8 to 12 dpi which maintained stable ASE rates (11.1–13.2%). Early infection at 4 dpi had the fewest (1.2%) ASE pairs, potentially due to the low fungal biomass sampled. To reduce noise in the subsequent ASE analyses, we applied a |LFC| threshold of two to define the ASE set (Diff2 and Diff4), while all other categories were defined as non-ASE, for the remainder of this study. Intersection analysis identified 678 allele pairs (8.3%) that were assigned to Diff2 or Diff4 ASE in at least one condition (Supplemental Fig. S13).

Given that pathogenicity-related effectors typically undergo diversifying selection (Sperschneider et al. 2014; Stukenbrock and McDonald 2009), we hypothesised that ASE might be an additional pathway to drive virulence dynamics. We therefore asked whether secretome genes, including putative effectors, were overrepresented in each condition-specific ASE versus non-ASE set compared to those of evolutionarily conserved BUSCOs. A Fisher’s exact test (FDR < 5%) for every condition consistently revealed significant ASE enrichment for secretome genes relative to BUSCOs as reflected by odd ratios greater than one (Fig. 6C; contingency tables in Supplemental Table S13).

We explored several epigenetic and genetic factors that potentially underlie the ASE of secretome genes. We compared the CpG methylation density between secretome alleles along their gene bodies (start to stop codon) and 2 kb flanking sequences to include proximal *cis* elements such as promoters. We observed significant methylation differences around the start codon between ASE alleles across all *in planta* infection conditions when compared to non-ASE counterparts (Fig. 6D, Supplemental Fig. S14). Higher-expressed alleles were found to be hypomethylated at the start codon windows (1.1–2.5% CpGs methylated), compared to those of their lower-expression counterparts which were frequently more heavily methylated (19.2–36.1% CpGs methylated). Non-ASE secretome genes showed no methylation difference between alleles regardless of their expression levels. These data suggest that asymmetrical methylation may be involved in secretome ASE.

Lastly, we investigated if SV or other genetic factors might be linked to methylation differences across ASE secretome genes. We compared TE occupancy in 5’ and 3’ flanking regions of ASE genes with the expectation that fungal TEs are typically heavily methylated (Bewick et al. 2019), but no significant difference was found between higher- and lower- expression alleles (Mann-Whitney U tests; Supplemental Fig. S15). Permutation tests also revealed depletions of ASE events within and nearby whole-genome SVs (Fig. 5C). Examination of allele sequence divergence showed no correlation between sequence divergence and expression differences of ASE secretome genes, except a weak positive correlation detected at 8 dpi (Spearman’s Rank-Order correlation tests; Supplemental Fig. S16). Hence it is currently unclear what drives ASE in *Pst*104E.

## Discussion

Recent advances in long-read sequencing technologies have made T2T haplotype-phased genome assemblies the new gold standard for eukaryotes, including di- and heterokaryotic fungi such as rust fungi. Here we report a fully nuclear-phased T2T genome assembly for *Pst,* the first reconstructed using high-accuracy ONT duplex sequencing. We show that our ONT-only T2T genome assembly is of comparable or superior quality to recently published PacBio HiFi-based haplotype-resolved T2T assemblies of other di- and heterokaryotic fungi (Wang et al. 2024; Sperschneider et al. 2023; Henningsen et al. 2024). With the complete resolution of both nuclear haplotypes in our *Pst* assembly, we were able to uncover novel insights into the impact of dikaryotism on the genome biology of a long-term asexual clonal isolate of this fungus.

Our detailed analysis of *Pst* centromeres shows that they adopt the classic Rabl configuration with clustering of heterochromatic centromeres (Torres et al. 2023; Xia et al. 2022). This arrangement allowed us to identify large regional centromeres from Hi-C contact hotspots that coincide with hypermethylated, gene-poor genomic signatures. Each centromere has a single potential kinetochore attachment site marked by a hypomethylated pocket known as the CDR (Logsdon and Eichler 2022; Logsdon et al. 2024; Mastrorosa et al. 2024), within otherwise fully CpG-methylated centromeres without signs of elevated AT content. Direct experimental evidence from CENP-A (CenH3) chromatin immunoprecipitation sequencing will be required to validate centromere localisation. Our comparative inter-haplotype analysis revealed that *Pst* centromeres are highly variable in length and sequence, lacking characteristic motifs that define all centromeres.

Remarkably, in most cases the inferred kinetochore site appears to be consistently positioned in homologous chromosomes even considering highly divergent homologous centromeres; and where shifts occur, we did not detect an associated sequence presence/absence pattern. These findings point to the conclusion that formation of *Pst* centromeres is a sequence-independent process, consistent with observations in many other fungal regional centromeres (Cissé et al. 2024; Schotanus et al. 2015; Roy and Sanyal 2011; Smith et al. 2012; Sperschneider et al. 2021). Most but not all *Pst* centromeres are enriched for LTR retrotransposons, especially the Ty3/Gypsy superfamily. LTR-rich centromeres have also been reported for other pathogenic fungi with large regional type centromeres, such as the closely related stem rust fungus *P. graminis* f. sp. *tritici* (Sperschneider et al. 2021), a human pathogenic yeast *Cryptococcus neoformans* (Yadav et al. 2018b) and an ascomycetous phytopathogen *Verticillium dahliae* (Seidl et al. 2020), but direct roles for LTR retrotransposons in centromere establishment remain unclear. In *C. neoformans*, the RNAi machinery and DNA methylation have been proposed as key epigenetic drivers for centromere identity via suppressing transposition and deleterious recombination among centromeric LTRs (Yadav et al. 2018b). However, the fact that the correlation with LTR enrichment does not hold in some *Pst* centromeres supports the idea that the LTR sequence itself does not define centromeres (Lynch et al. 2010; Guin et al. 2020). Rather, LTRs might be preferentially inserted due to their high proliferative potential which may, in turn, promote RNAi-directed silencing to reinforce centromere formation (Balzano and Giunta 2020). Further studies will be required to elucidate such links in *Pst*.

The dikaryotic configuration of *Pst* prompted us to explore its effect on intraspecific rDNA dynamics under strict clonality. Generally, rDNA tandem repeats are thought to undergo concerted evolution towards sequence homogenisation via repeated homologous recombination, namely unequal crossovers and gene conversion (Garcia et al. 2024; Symonová 2019; Mullis et al. 2020). Because *Pst*104E has genetically distinct nuclei, we hypothesised that its rDNA variants may persist or emerge within individual nuclei in the absence of meiotic exchange. Our results show that each nucleus of *Pst*104E carries a unique array with over 200 repeats, predominated by an rDNA sequence subtype that harbours nucleus-specific variations within both IGS (intergenic spacer) regions. A low proportion of these repeats have diversified through accumulating point mutations, and some appeared to have become fixed. We speculate that such nucleus-specific rDNA subtype homogeneity might be the consequence of compartmentalised concerted evolution due to the individual inheritance of each nucleus over prolonged clonal history. The intra-array diversification, however, signifies a relaxation of concerted evolution leading to incomplete homogenisation within each nucleus (Wang et al. 2023; Xu et al. 2017). A possible explanation for this could be reliance on limited non-meiotic homologous recombination (e.g. intrachromosomal or between sister chromatids) that can rapidly purge the newly spreading variants (Paloi et al. 2022). This parallels previous observations from multinucleate arbuscular mycorrhizal fungi which, unlike *Pst*, are homokaryotic (i.e. have genetically uniform nuclei) (Serghi et al. 2021), but display extensive rDNA heterogeneity within each nucleus, consistent with their ancient clonality (Lin et al. 2014; Pawlowska and Taylor 2004). In future, it will be interesting to survey intraspecific rDNA variations in rust isolates that have arisen via recent sexual recombination (Wang et al. 2022; Heitman et al. 2013; Wallen and Perlin 2018), where we expect that divergent rDNA subtypes will be more evenly distributed among nuclei. Dikaryotism therefore presents an excellent opportunity for understanding the rates and dynamics of concerted evolution in fungi.

A fundamental question in genome biology is whether ASE could have functional consequences that lead to phenotypic variability (Cleary and Seoighe 2021; St. Pierre et al. 2022). Here, we show that ASE is pervasive in *Pst* and appears to be inversely related to gene body methylation, where lower-expressed alleles display higher levels of CpG methylation. In *Pst*104E, ASE is overrepresented in secretome genes including putative effectors that could be involved in host pathogenesis. Therefore, ASE might present a novel regulatory mechanism to generate effector diversity beyond simple protein sequence variations (Chen et al. 2017; Salcedo et al. 2017; Ortiz et al. 2022). Expression level polymorphisms between recognised and non-recognised effector alleles may also explain virulence switching. For example, in the stem rust fungus *P. graminis* f. sp. *tritici*, a virulence allele of *AvrSr27* was expressed at a much lower level than its avirulence allele counterpart; yet when over- expressed *in planta*, recognition still took place (Upadhyaya et al. 2021). Non-recognition is therefore due to low expression rather than an inability of the host and pathogen proteins to interact. Similar observations have been recently made in a common rust disease of maize caused by *P. sorghi*. A lowly-expressed virulence allele of *AvrRp1-*D differing by only one amino acid from its avirulence counterpart was recognised by the cognate resistance gene *Rp1-D* after co-overexpression *in planta* (Kim et al. 2024). Future studies will inform whether the ASE of these *P. graminis* f. sp. *tritici* and *P. sorghi* effector genes of is linked to changes in gene body methylation.

The role of epigenetic regulation of *Avr* gene expression was previously highlighted in the soybean pathogen *Phytophthora sojae*, where natural silencing of the avirulence gene *Avr3a* resulted in gain-of-virulence (Qutob et al. 2013; Hale et al. 2023). Such epigenetics-mediated ASE may offer a reversible means to selectively silence or “archive” avirulence alleles to escape immune recognition while retaining the unmutated gene.

One important limitation of our study is that our DNA methylation data only reflect dormancy. Chromatin remodelling via histone modifications (which are highly correlated with DNA methylation in fungi) (Rose and Klose 2014; He et al. 2020) can de-repress effector expression in *Zymoseptoria tritici* during pathogenesis (Meile et al. 2020). To extend our observations, we need to DNA methylation and histone modification data sampled from *Pst* during pathogenic growth *in planta* to validate if chromatin dynamism underpins ASE *in planta*.

For now, the plot thickens; Flor’s classic ‘gene-for-gene’ hypothesis (Flor 1942) that has shaped our understanding of plant resistance to pathogens since the 1940’s might be an oversimplification, and we may have to consider that differences in effector gene expression underly disease outcomes in the field.

## Methods

### DNA extraction and long-read sequencing

High-molecular-weight genomic DNA was extracted from fresh Pst104E urediniospores as described previously (Schwessinger and Rathjen 2017) and size-selected for at 15kb using the BluePippin (Sage Science). Nanopore sequencing library was prepared using the Ligation Sequencing Kit v14 (SQK-LSK114), then sequenced on a PromethION sequencer, each library on a single R10.4.1 flowcell at 260 bps translocation speed. Sequencing was performed at the Biomolecular Resource Facility (BRF) at The Australian National University. Simplex reads were basecalled from raw signals with dorado v0.2.1 using the super accuracy (SUP) model “dna_r10.4.1_e8.2_260bps_sup@v4.1.0”. Duplex reads were paired with duplex-tools v0.3.1, then basecalled using the stereo model “dna_r10.4.1_e8.2_4khz_stereo@v1.1”. Seqkit v2.6.1 (Shen et al. 2016) was used to confirm the sequencing quality of each library’s output before concatenating them. NanoFilt v2.8.0 was used to trim and filter duplex reads (-q 30 -l 10000 --headcrop 75 -- tailcrop 75), as well as simplex reads (-q 10 -l 40000). To address chimera, we performed all-versus-all duplex read alignment with minimap2 v2.26 (-x ava-ont) (Li 2018), followed by yacrd and fpa (Marijon et al. 2020) to conservatively split chimeric reads at zero coverage regions.

### Hi-C sequencing

Spore samples were cross-linked with 1% formaldehyde, quenched with 1% glycine, washed twice with 1x PBS and ground to fine powder using a Qiagen TissueLyser I. Tissue was sent to Phase Genomics (Seattle, WA, USA) for Hi-C library preparation (Proximo Hi-C (Fungal) Kit KT6040 Protocol Version 4.0 (February 2021), restriction enzymes used: DpnII, HinFI, MseI, DdeI) and sequencing. Sequencing (150 bp paired-end) was performed on an Illumina NovaSeq 6000 flowcell. Hi-C library was quality controlled by quantifying valid interactions in the resulting sequencing data with HiC-Pro v3.1.0 (Servant et al. 2015).

### Long-read cDNA library preparation and sequencing

Infected leaves and urediniospores were ground to fine powder using a Qiagen TissueLyser II (25 Hz, 1 min). 1 mL of TRIzol (Invitrogen, 15596018) was added to each sample and RNA was extracted with the Zymo Research Direct-zol RNA Miniprep Plus Kit (Zymo Research, R2070), including DNase I treatment. RNA concentration was determined with the Qubit RNA BR Kit (Invitrogen, Q10211), RNA integrity and quality was assessed with the Qubit RNA IQ Assay Kit (Invitrogen, Q33221) and by agarose gel electrophoresis.

50 ug total RNA were used for poly-A enrichment with Dynabeads Oligo(dT)_25_ (Invitrogen, 61005) according to manufacturer’s instructions for purifying mRNA from total RNA. First strand cDNA synthesis was based on a modified version of the Oxford Nanopore “Direct cDNA Sequencing V14 with SQK-LSK114” protocol. Second strand synthesis and RNA degradation were based on a modified NEB protocol (https://www.neb.com/en-au/protocols/2019/05/09/2nd-strand-cdna-synthesis-protocol-using-the-template-switching-rt-enzyme-mix). See Supplementary Notes for detailed descriptions. Barcoding and library preparation of samples were performed with the Oxford Nanopore technologies (ONT) Native Barcoding Kit 96 (SQK-NBD114.96) using approximately 100 ng cDNA as input. Barcoded samples were combined in different pools for ONT sequencing using three FLO-PRO114M flowcells. Sequencing was performed at the Biomolecular Resource Facility (BRF) at The Australian National University.

### Genome assembly and Hi-C scaffolding

We used Verkko v1.3.1 (Rautiainen et al. 2023) to assemble 32x duplex (--hifi) and 117x simplex (-- nano) reads following developers’ recommendation for ONT-only assembly. To incorporate Hi-C, we ran the “gfase_wrapper.sh” script from an early version of Verkko’s Hi-C phasing pipeline (https://github.com/Dmitry-Antipov/verkkohic), which employs bwa v0.7.17-r1188 (Li and Durbin 2009) to map Hi-C data against the raw assembly, and GFAse (Lorig-Roach et al. 2023) to perform haplotype phasing. The raw assembly was aligned to the previously published *Pst*134E assembly (Schwessinger et al.) using D-GENIES (Cabanettes and Klopp 2018) to identify chromosome-scale contigs and homologous pairs. Contaminant and mitochondrial contigs were identified using local BLAST v2.14.0 search against the NCBI’s nt database (blastn -perc_identity 75 -evalue 1e-5) for removal. Contigs with mean window coverage lower than 5x were discarded. Hi-C scaffolding was then performed using Juicer v2.0 (Durand et al. 2016b) and 3D-DNA v180114 (Dudchenko et al. 2017), followed by gap filling and telomere recovery as described in the Supplementary Notes. Hi-C heatmap was visualised in Juicebox (Durand et al. 2016a). The karyotype plot was plotted using karyoploteR (Gel and Serra 2017).

### Assembly evaluation

Assembly quality was evaluated for the dikaryotic assembly and each of the phased haplotypes. Basic assembly statistics, such as N/L50 and GC content, were generated using Seqkit v2.6.1 (Shen et al. 2016). BUSCO v5.5.0 (Simão et al. 2015) was launched in nucleotide mode to count core genes present in the basidiomycota_odb10 database (downloaded on 8 January 2024). To obtain the k-mer-based consensus QV score, 31-mers were generated from duplex reads using meryl v1.4 (Rhie et al. 2020), followed by Merqury v1.3 (Rhie et al. 2020) to evaluate concordance with the assembly to compute QV. LAI score was obtained by running LTR_retriever (Ou et al. 2018; Ou and Jiang 2018) on the suggested settings to identify LTR retrotransposons and assess contiguity. Base-level discrepancies were identified by calling SNPs from duplex read alignment using bcftools v1.19.1 *mpileup* and *call* (-mv to include multiallelic sites) (Danecek et al. 2021) functions. We also launched CRAQ v1.0.9 (Li et al. 2023b) on both Illumina and duplex alignments to find structural discrepancies. The output R/S-AQI scores are inferred from regional and structural errors from clipped alignments that may indicate misjoins. Any identified discrepancies were flagged for further examination and, where appropriate, manually curated using Flye v2.9.1 (Kolmogorov et al. 2019) to perform local assembly of nearby UL reads. To evaluate phasing quality, HiC-Pro was used to generate contact matrices from Hi-C alignments with a minimum MAPQ of 20. The contact matrices were then analysed using python scripts from https://github.com/RunpengLuo/HiC-Analysis/tree/main (Luo 2024) to quantify *cis-* and *trans-*chromosome Hi-C contacts. Rolling means over 20 Kbp windows were plotted onto a circos plot to spot potential phase switch between the haplotype assemblies.

### TE annotation

Each haplotype genome of *Pst*104E was annotated separately. TEs were predicted *de novo* using the REPET v3.0 pipeline, which consists of TEdenovo (Flutre et al. 2011) and TEannot (Quesneville et al. 2005). First, TEdenovo was run on default settings to detect repeats based on Repbase v27.06 (nt and aa databases), using the hidden Markov model profile bank from Pfam v35.0 (Finn et al. 2013) and Gypsy Database v2.0 (Lloréns et al. 2007) (“ProfilesBankForREPET_Pfam35.0_GypsyDB_2022.hmm”). Then, following REPET authors’ recommendations for better annotation quality, two rounds of TEannot were performed. The first round was to define the total TE consensus library without feature detection and classification (steps 1, 2, 3 and 7). Consensuses that have at least two copies and one full-length copy (i.e. matching fragments aligned with >95% of the consensus length) were retained as validated TEs, which were then annotated in the second TEannot run (steps 1–5, 7 and 8). PASTEC (Hoede et al. 2014) classifications and statistics for each TE consensus were obtained from output files with the “.classif” and “.annotStatsPerTE.tab” extensions, respectively. Bedtools v.2.30.0 *maskfasta* (Quinlan and Hall 2010) was used to soft-mask the annotated TEs in the genome assembly.

### Transcriptome assembly

Prior to gene annotation, we independently processed the previously published Illumina RNA-seq (Schwessinger et al. 2018) and our ONT long-read cDNA datasets to generate transcript evidence. Illumina RNA-seq was filtered and trimmed with fastp v0.23.4 (--detect_adapter_for_pe --cut_right -- correction) (Chen et al. 2018). Processed reads were mapped to the dikaryotic assembly using HISAT2 v2.2.1 (--max-intronlen 3000 --dta) (Zhang et al. 2021). Aligned and paired reads (-F4 -f2) were partitioned into haplotype sets using samtools v1.18 (Li et al. 2009) and merged across replicates (n=3) per condition. Transcriptome assembly was then performed for each condition. For *Pst*104E, CYR32 and *Pst*87/66 samples, we used StringTie2 (v2.2.1) (Kovaka et al. 2019) to infer transcripts from splice alignments, with -s2 and -m50 applied to ensure capturing of shorter single-exon transcripts. Additionally for *Pst*104E, we assembled transcripts using reference-guided Trinity v2.9.1 (--genome_guided_max_intron 3000 --jaccard_clip) (Grabherr et al. 2011).

For the *Pst*104E ONT cDNA dataset, reads were trimmed with Porechop_ABI v0.5.0 (Bonenfant et al. 2023), then aligned to the dikaryotic assembly using minimap2 v2.26 in splice-aware mode (-ax splice -ub -G 3000 --secondary=no). Aligned reads were partitioned into haplotype sets and merged across replicates (n=4) per condition. To identify transcript structures from noisy long reads reference-guided and annotation-free, we employed two recently published tools: StringTie2 (-L -s2 -m50) for better single-exon transcript discovery, and ESPRESSO v1.4.0 (ESPRESSO_S.pl -Q0) (Gao et al. 2023) for improved splice site detection. All the transcript GTF annotations were merged across all *Pst* samples using StringTie2 (--merge). For *Pst*104E, we also extracted transcriptome FASTA sequences from the annotations using Gffread v0.12.7 (Pertea and Pertea 2020), and concatenated them to the Trinity assemblies.

### Gene annotation

Gene annotation was carried out separately for each haplotype. Funannotate v1.8.15 (Jonathan and Jason 2023) was launched to train PASA on all preassembled *Pst*104E transcripts from ESPRESSO, StringTie2 and Trinity. CodingQuarry-PM (pathogen mode) v2.0 (Testa et al. 2015) was run on to predict genes from the merged transcript annotations. Both standard and dubious outputs were combined into a single CodingQuarry-PM gene set. Next, we executed funannotate *predict* (-- optimize_augustus --ploidy 1 --repeats2evm) using inputs as followed: *Pst*104E transcriptome assemblies (--transcript_evidence); transcript alignments (--rna_bam); transcript annotations from ESPRESSO and StringTie2 (--transcript_alignments), trained PASA (--pasa_gff), CodingQuarry-PM (--other_gff); as well as protein evidence from UniProtKB/Swiss-Prot (release 2023_04) (The UniProt Consortium 2023) and the previously published *Pst*104E proteome (Pst_104E_v13_ph_ctg.protein.fa) (Schwessinger et al. 2018). All evidence types were parsed to EvidenceModeler v1.1.1 (Haas et al. 2008) to produce weighted consensus gene structures.

Evidence weights were configured as “augustus:4 hiq:6 genemark:1 pasa:10 codingquarry:0 snap:1 glimmerhmm:1 proteins:6 transcripts:6”, and “--other_gff:10” for CodingQuarry-PM input. Untranslated regions were inferred from long-read alignments using InGenAnnot *utr_refine*.

For functional annotation, we started by predicting secretome and effector genes using SignalP v6.0 (--organism eukarya --mode slow) (Teufel et al. 2022) and InGenAnnot v0.0.11 *effector_predictor* (Lapalu et al. 2023). InGenAnnot *rescue_effectors* was employed to find potential effector genes missed in the unannotated transcripts. To filter false positives, we used Phobius v1.01 (Käll et al. 2004) and TMHMM v2.0 (Krogh et al. 2001) to confirm the absence of transmembrane domain outside of the N-terminal signal peptide region; only hits predicted by both tools were removed as they are more likely to be biologically accurate. Seconary metabolite biosynthesis gene clusters were identified using antiSMASH v6.1.1 (Medema et al. 2011). InterProScan v5.64-96.0 (Blum et al. 2021; Jones et al. 2014) was run locally to predict protein functions using its defaulted member databases. All results were parsed to funannotate *annotate* to integrate functional annotations to the predicted genes. BUSCO completeness of the annotated genes was assessed in protein mode.

### Differential gene expression analysis

Transcript abundance was quantified from *Pst*104E ONT cDNA alignments using bambu v3.4.1 (Chen et al. 2023) with isoform discovery mode disabled. The resulting count matrices were imported to DESeq2 v1.38.3 (Love et al. 2014) for analysis. To assess data quality, PCA was performed on variance-stabilising transformed read counts to visualise sample clustering. DESeq2 differential expression analysis was then conducted using default settings to identify genes differentially expressed *in planta* compared to UG. DESeq2 applies the Wald test to determine the statistical significance of expression LFC for each gene, with p-values adjusted for multiple testing using the Benjamini and Hochberg method. We considered genes with an adjusted p-value<0.05 and LFC≥2 to be upregulated. Secretome genes upregulated during 4 or 6 dpi were identified as our preliminary *Avr* effector candidates.

### Centromere inference and analysis

Centromere locations were first estimated from the genome-wide Hi-C heatmap in Juicebox by identifying strong inter-chromosomal bowtie-like contact signals indicative of centromere-to-centromere interactions. To assess their methylation status, we called 5mCG modifications from a 400 bps ONT genomic dataset of *Pst*104E using dorado’s SUP model “dna_r10.4.1_e8.2_400bps_sup@v4.1.0”. Reads with methylation calls were mapped to the *Pst*104E assembly with minimap2 (-ax map-ont --secondary=no). Modkit v0.2.5 *pileup* (https://github.com/nanoporetech/modkit) was used to count modified CpGs in the reference using the aligned reads (--edge-filter 75 --combine-strands --bedgraph). This generates a bedGraph file that reports the fraction of reads exhibiting cytosine methylation at each reference CpG. We calculated mean methylation fractions over 500 bp windows and plotted them along each chromosome to confirm overlap between methylation peaks and the Hi-C bowtie signals, which we inferred as centromeres. To analyse their synteny, pairwise alignment between homologous centromeres was performed using MUMmer’s v4.0.0rc1 NUCmer tool (Marçais et al. 2018) with default settings. Alignment blocks longer than 100 bp with minimum sequence identity of 90% were retained and plotted in Dot (Sommer 2021).

We determined the locations and sizes of CDRs by visually selecting the largest hypomethylation region as this single pattern consistently appeared throughout all the centromeres. Relative CDR positions were calculated as the midpoint coordinate divided by centromere length, then compared between haplotypes to detect CDR shifts. To investigate sequence composition at shifted CDRs in greater detail, high-resolution alignment dotplots of homologous centromeres were generated using Gepard (Krumsiek et al. 2007). The associated TEs were inspected in Geneious and visualised as feature tracks using pyGenomeViz (Shimoyama 2024).

### Centromeric TE enrichment analysis

Centromeric TE enrichment was analysed using a custom script employing permutation tests. The script takes in centromere and TE annotations, then applies bedtools *intersect* (-f 0.5) to identify TEs located in centromeric and non-centromeric regions along with their classifications. TE locations are then randomly shuffled along each chromosome in 5,000 permutations using bedtools *shuffle*. For every permutation, the coverage difference of a given TE superfamily between the centromeric and non-centromeric region is calculated. All permuted data generates a null distribution, enabling a two-tailed test for the statistical significance of centromeric enrichment or depletion per TE superfamily.

P-values, defined as the proportion of permuted results equal to or more extreme than the observed, were adjusted for multiple testing using <5% FDR.

### Ribosomal DNA analysis

rDNA arrays were located using the whole-genome BLAST results as described above. We defined the canonical rDNA unit by integrating evidence from homology alignment and rRNA reads from the ONT cDNA library. Reference sequences of ITS, 18S and 5S regions from *Puccinia* species were retrieved from public databases including Gold Standard (Eenjes et al. 2022), EukRibo (Berney et al. 2022) and 5SrRNAdb (Szymanski et al. 2016), and aligned to the rDNA arrays to distinguish conserved and variable elements. Long rRNA reads were then used to refine rRNA gene boundaries. To suppress multimapping, all rDNA sequences were removed from the assembly, with a single rDNA repeat (starting from 18S) added as an extra “scaffold”. All UG cDNA reads were mapped to the edited assembly with minimap2 in splice-aware mode, then visualised in IGV to reveal transcribed elements and confirm the canonical rDNA unit.

To capture rDNA sequence variations, duplex reads were aligned to the same edited assembly with minimap2, revealing two dominating subtypes. Reads were then mapped to these subtypes to identify low-frequency SNPs, which were called using bam-readcount v1.0.1 (Khanna et al. 2022) and parsed into a SNP information table with the included “parse_brc.py” script. SNPs were filtered with a 30x alternate base count threshold, as this reflects the lowest duplex read depth expected for single-copy subtypes based on the haploid genome coverage. SNPs detected at homopolymers were also excluded. SNP combinations were manually examined in read alignments, allowing us to reconstruct twelve low-frequency rDNA subtypes. Multiple sequence alignment among all subtypes was conducted using MAFFT v7.490 (Katoh et al. 2002; Katoh and Standley 2013). Inspired by (Sharma et al. 2022), the copy number of each rDNA subtype was estimated by normalising duplex and Illumina read depths (or SNP depths for low-frequency subtypes) to the mode per-base depth values of the whole genome, rather than the mean-based approach to minimise skewness.

The nuclear specificity of the two dominant rDNA subtypes was tested by analysing rDNA Hi-C reads with a k-mer approach. Unique 31-mers for each subtype were identified using UniqueKMER (Chen et al. 2020), with their reverse complements added via seqkit (Shen et al. 2016). Subtype-specific 31-mers were used to tag rDNA Hi-C reads with UNIX grep command. Using their read identifiers, paired Hi-C mates were fetched from the corresponding R1/R2 read file. Mates were mapped to the dikaryotic assembly with bwa-mem2 (Vasimuddin et al. 2019), and processed with bedtools *bamtobed* to extract mapping locations and MAPQ scores for plotting. This was repeated on subtype-unspecific 31-mers as control (note if a read is tagged by both subtype-specific and unspecific *k*-mers, it is defined as subtype-specific).

### Synteny, structural rearrangements and variations detection

To assess interhaplotype synteny, we performed whole-genome alignment between haplotypes A (reference) and B (query) using NUCmer (--maxmatch -l 200 -b 500 -c 500). Alignment blocks with <90% identity were filtered out using MUMmer’s delta-filter, and the resulting delta file was converted to alignment coordinates using show-coords (-THrd) for downstream SV calling. SyRI (Goel et al. 2019) was launched on default settings to annotate structural rearrangements (inversions, translocations and duplications), syntenic regions, and unaligned (sequences absent in one haplotype due to indels or excessive sequence divergence) regions. Visualisation of synteny and SVs between homologous chromosomes were generated by plotsr (Goel and Schneeberger 2022). To analyse features within and nearby SVs, bedtools *slop* was used to extend SV coordinates by 2 kbp in both directions. Enrichment or depletion for genomic features, including TEs, genes, and specific subsets like secretome-only and ASE-only genes, was statistically assessed through two-tailed permutation tests as described above.

### Identification of hemizygous and heterozygous genes

Protein sequences from all annotated genes were analysed using Proteinortho v6.3.1 (Lechner et al. 2011) with the -synteny flag to detect homologs between haplotypes. Genes lacking a hit on the alternative haplotype were considered hemizygous candidates. These were further filtered via reciprocal BLASTp to ensure the absence of alleles. Candidates with a high-quality hit that had >70% identity and >70% query and subject coverage were omitted, producing the final hemizygous gene list, which we intersected with upregulated secretome genes for high-priority *Avr* candidates.

For heterozygous genes, we began by identifying one-to-one gene pairs from Proteinortho results. Using a script adapted from “dN_dS_Pst134E.ipynb” (https://github.com/ZhenyanLuo/codes-used-for-mating-type (Luo et al. 2024), MUSCLE v3.8.31 (Edgar 2004) was run on default settings to perform codon-aware alignments between each gene pair. Synonymous (d_S_) and non-synonymous (d_N_) divergence values were then calculated using PAL2NAL v14 (Yang 2007), along with CDS and protein Levenshtein distances generated by editdistance v0.6.2. A gene pair was determined to be heterozygous biallelic if either its d_S_ or d_N_ value was greater than zero.

### Allele-specific expression analysis

Having the paired allele information, we reformatted the bambu per-gene cDNA count matrices for testing condition-specific ASE. Each row represents an allele pair, while counts for one allele and the other across all samples appear as subsequent columns (48 in total). With haplotype A alleles set as the reference, differential expression between allele pairs was analysed for each condition using DESeq2 on default settings. The resulting |LFC| and adjusted p-values were used to categorise the allele pairs into different ASE status, as detailed in the Results. To assess nuclear dominance, allele read counts were normalised using DESeq2’s median of ratio method and transformed into expression levels as log_10_(median of ratio+1). Mean allele expression levels were calculated by averaging across replicates for haplotype comparisons.

To test for the overrepresentation of secretome genes in ASE set (Diff2 and Diff4) compared to BUSCOs, six 2x2 contingency tables were constructed using counts of ASE or non-ASE secretome and BUSCO genes for each condition. Two-sided Fisher’s exact tests of independence were performed using SciPy v1.9.3 to compute the odds ratio and p-values, corrected to <5% FDR.

CpG methylation differences between ASE and non-ASE secretome alleles were analysed using a custom python script. Gene body regions were defined based on the start and stop codon coordinates from CDS GFF3 annotations, then extended outwards by 2 kbp to include upstream and downstream flanking regions, generating three BED files per gene. Each region was divided into 20 equally proportioned bins, totaling 60 bins per gene. Mean methylation density was calculated per bin and averaged across higher- and lower-expression alleles for each condition.

Bootstrapping with 1,000 replicates was used to compute 95% confidence intervals.

To investigate TE occupancy surrounding the ASE genes, overlapping TE fragments were merged with bedtools *merge*, followed by bedtools *intersect* (-wo) to report the number of bases covered by TEs within each 5’ and 3’ flanking region.

## Data access

All sequencing data generated for this paper has been deposited in the Sequence Read Archive registered under BioProject accession number PRJNA1195871. All custom scripts and codes for analyses and generating the figures are available on GitHub: https://github.com/ritatam/Pst104EGenomeBiology. Genome assembly, gene annotations and TE annotations are available on Zenodo (doi: 10.5281/zenodo.14398118).

## Acknowledgements

1. R. T. was supported by a Grains Research and Development Corporation Graduate Research Scholarship. This work was supported by an Australian Research Council Future Fellowship to B.S. (FT180100024) and a Discovery Project Grant (DP230100941) to J.P.R. and B.S. This work was supported by computational resources provided by the Australian Government through the National Computational Infrastructure (NCI) under the ANU Merit Allocation Scheme. We would like to acknowledge the contribution of the Plant Pathogen ‘Omics Initiative consortium in the generation of data used in this publication. The Initiative is supported by funding from Bioplatforms Australia, enabled by the Commonwealth Government National Collaborative Research Infrastructure Strategy (NCRIS).

