## Supplemental Material for "Long-read genomics reveal extensive nuclear-specific evolution and allele-specific expression in a dikaryotic fungus"

**Supplemental Table legends**

**Supplemental Table S1.** Summary statistics of the raw Verkko assembly of *Pst*104E.

**Supplemental Table S2.** Summary statistics of the time-course ONT direct cDNA sequencing dataset for *Pst*104E. This includes number of reads, read length, percentage of read quality above Q15, number and percentage of reads mapped to the *Pst*104E assembly, etc.

**Supplemental Table S3.** Metadata and citations for all public Illumina RNA-seq datasets for three different *Pst* isolates (*Pst*104E, *Pst*87/66 and CYR32) used for transcriptome assembly and gene annotations in this study.

**Supplemental Table S4.** Information on the 89 hemizygous *Avr* candidates identified via DESeq2 differential gene expression analysis based on ONT cDNA transcript abundance. Columns are listed in the following order: gene identifier, expression log_2_ fold change in all *in planta* conditions (4 to 12 days post infection) relative to the ungerminated urediniospores (dormancy) condition, coding region lengths, and the associated functional annotations.

**Supplemental Table S5.** Summary statistics of transposable elements (TEs) identified and classified for each of the nuclear haplotype of the *Pst*104E genome assembly using the REPET pipeline.

**Supplemental Table S6.** Coordinates and length of the inferred centromeres and the centromere dip region (CDR). The CDR midpoint was used to calculate its relative position by dividing its coordinate by the centromere length, then compared to the relative CDR position on its homologous centromere.

**Supplemental Table S7.** Permutation test results for centromeric TE superfamily enrichment or depletion, conducted for each chromosome. The test statistic is the TE coverage difference between centromeric and non-centromeric region. TE locations are randomly shuffled along each chromosome in 5,000 permutations to generate a null distribution, enabling a two-tailed test for the statistical significance of centromeric enrichment or depletion per TE superfamily. P-values, defined as the proportion of permuted results equal to or more extreme than the observed, were adjusted for multiple testing using <5% FDR.

**Supplemental Table S8.** Information on the rarer rDNA subtype variants (#1.1-1.9 and #2.1-2.3) identified via calling low-frequency SNPs from ONT duplex alignment against the two dominant rDNA subtypes #1 and #2. The SNP positions and the overlapping rRNA gene annotations are listed in the top two rows. Point mutations were scored as “reference>alternate(SNP depth)”. SNP combinations for each subtype variant was visually determined in the IGV alignment. SNP depth was used to estimate their copy number by normalising it against the mode value of genome-wide per-base coverage depth.

**Supplemental Table S9.** SyRI summary statistics of the structural variations (SVs), syntenic and unaligned (non-syntenic) regions identified between haplotypes.

**Supplemental Table S10.** Permutation test results for the enrichment or depletion of different genomic features (TEs, genes, secretome/effector genes and allele-specific expressed genes) within different SV types conducted for whole genome. Statistical significance was assessed through two-tailed permutation tests, as described above for centromeric TE analysis.

**Supplemental Table S11.** Proteinortho results and divergence values for one-to-one allele pairs identified between *Pst*104E haplotypes A and B. Identified hits from the Proteinortho output table were filtered based on e-value < 0.05. Synonymous (dS) and non-synonymous (dN) divergence values along with CDS and protein Levenshtein distances were then calculated. A gene pair was determined to be heterozygous biallelic if either its dS or dN value was greater than zero.

**Supplemental Table S12.** DESeq2 results for allele-specific expression (ASE) analysis conducted on the heterozygous biallelic gene pairs, with haplotype A alleles set as the reference. The resulting |LFC| and adjusted p-values (padj) were used to categorise the allele pairs into different ASE status, as detailed in the Results.

**Supplemental Table S13.** Two-by-two contingency table of counts of ASE versus non-ASE allele pairs that have at least one allele annotated as secretome gene or BUSCO. Fisher's exact test (two-sided) was applied to obtain the odds ratio and Fisher's exact p-value, adjusted for multiple testing using <5% FDR. UG: ungerminated spores; dpi: days post infection.

**Supplemental Notes**

**Long-read cDNA library preparation and sequencing**

Infected leaves and urediniospores were ground to fine powder in 2 mL tubes containing metal beads with a Qiagen TissueLyser II (25 Hz, 1 min). 1 mL of TRIzol (Invitrogen, 15596018) was added to each sample and samples were stored at -80°C until RNA extraction. For RNA extraction, samples were incubated in TRIzol (Invitrogen) for 10 min at room temperature and centrifuged at 10,000 g for 5 min. Supernatant was transferred to a new tube and an equal amount of 100 % EtOH added. The Zymo Research Direct-zol RNA Miniprep Plus Kit (Zymo Research, R2070) was used for isolation of RNA, including DNase I treatment. RNA concentration was determined with the Qubit RNA BR Kit (Invitrogen, Q10211), RNA integrity and quality was assessed with the Qubit RNA IQ Assay Kit (Invitrogen, Q33221) and by agarose gel electrophoresis.

50 ug total RNA were used for poly-A enrichment with Dynabeads Oligo(dT)_25_ (Invitrogen, 61005) according to manufacturer’s instructions for purifying mRNA from total RNA. Per sample, 80 µL Dynabeads Oligo(dT)_25_ were used and beads were reused for a second round of mRNA isolation for each sample to increase mRNA yield. First strand cDNA synthesis was based on a modified version of the Oxford Nanopore “Direct cDNA Sequencing V14 with SQK-LSK114” protocol. Approximately 150 ng of poly-A enriched RNA were used as input for 1^st^ strand synthesis including template switching to increase the number of full-length transcripts. First, oligo dT_23_ (/5phos/ ACTTGCCTGTCGCTCTATCTTCTTTTTTTTTTTTTTTTTTTTTTTVN, IDT, RNase free HPLC) primer binding was performed in the following reaction: 7.5 µL poly-A mRNA (~150 ng), 2.5 µL 2 µM oligo dT_23_ primer and 1 µL 10 mM dNTPs (NEB, N0447L) were incubated at 65°C for 5 min, then snap-cooled in a pre-chilled freezer block. First strand synthesis and template switching were performed by adding 4 µL 5X RT Buffer (Thermo Scientific, EP0751), 1 µL RNaseOUT (100 mM, Invitrogen, 10777019), 1 µL nuclease-free water and 2 µL 10 µM template switching oligo (/5phos/TTTCTGTTGGTGCTGATATTGCTrGrGrG, IDT, RNase free HPLC), incubation for 2 min at 42°C followed by addition of 1 µL (200 Units) Maxima H Minus Reverse Transcriptase (Thermo Scientific, EP0751). Reaction was performed at 42°C for 90 min, followed by heat-inactivation for 5 min at 85°C. Second strand synthesis and RNA degradation were based on a modified NEB protocol (<https://www.neb.com/en-au/protocols/2019/05/09/2nd-strand-cdna-synthesis-protocol-using-the-template-switching-rt-enzyme-mix> *add as reference*). The first strand synthesis product (20 µL) was mixed with 50 µL Q5® High-Fidelity 2X Master Mix (NEB, M0492L), 4 µL 10 µM PR2 primer (/5Phos/TT TCTGTTGGTGCTGATATTGC, IDT, HPLC), 5 µL (25 Units) RNase H (NEB, M0297L), 1 µL RNase Cocktail™ Enzyme Mix (Invitrogen, AM2286) and 20 µL nuclease-free water. The reaction was incubated at 37°C for 15 min to hydrolyse RNA, 95°C for 1 min for initial denaturation, 50°C for 1 min for primer annealing and 65°C for 15 min for second strand extension. The second strand synthesis product was cleaned up with 0.8x volume 2% SeraMag beads and eluted in 21 µL nuclease-free water. Barcoding of samples was performed with the Oxford Nanopore technologies (ONT) Native Barcoding Kit 96 **(SQK-NBD114.96) using approximately 100 ng cDNA as input. Barcoded samples were combined in different pools for sequencing on PromethION using three** FLO-PRO114M flowcells. Sequencing was performed at the Biomolecular Resource Facility at The Australian National University.

***Pst*104E assembly curation**

In the Verkko raw assembly of *Pst*104E, most chromosomes were directly resolved into full linear sequences as indicated by string graph in Supplemental Fig. S3. Out of the 36 total chromosomes, Verkko directly assembled 27 into gapless T2T contigs and one into T2T scaffold (chr13B; gap introduced at the repetitive rDNA array). Three chromosomes were broken into two to three contigs (chr6A, 8A and 13A, excluding rDNA fragments), which were joined into three additional chromosomal scaffolds using Hi-C linkage information as described in Methods. With the centromeres inferred from the Hi-C heatmaps, most chromosomes appeared to be acrocentric or submetacentric; we oriented them such that each begins from the shorter p-arm. We sorted the *Pst*104E chromosomes by the average length of each homologous pair starting from the longest to shortest.

We identified assembled telomeres by searching TTAGGG/CCCTAA motifs at the first and last 50bp of all scaffolds using “FindTelomeres.py” (<https://github.com/JanaSperschneider/FindTelomeres>) (Sperschneider 2024). Eight chromosomes had at least one telomere missing (chr2B, 5B, 6A, 8A, 10A, 11A, 13A and 16A), as seen in IGV with “show soft-clipped bases” option toggled on. To recover missing telomeres, we used the pre-release version of Teloclip (<https://github.com/Adamtaranto/teloclip>) (Taranto 2024) to extract UL reads aligned at contig ends and had soft-clipped overhangs harbouring at least two telomeric repeats, including the ~5 Kbp subtelomeric region for subsequent anchoring. These were locally assembled with Flye v2.9.1 (Kolmogorov et al. 2019), then stitched to the corresponding scaffold end. Only repeats supported by at least two reads were added. Reads were then mapped against the revised assembly to check if alignments correctly extended into the recovered ends.

For curation, we performed coverage check by separately mapping duplex and simplex reads back to the assembly with minimap2 (-ax map-ont) (Li 2018), then generated a genome-wide coverage plot via JVarkit’s *wgscoverageplotter* (Lindenbaum 2015). For better resolution, we used bedtools’ *makewindows* function (Quinlan and Hall 2010) to split contigs into 10kbp sliding windows with 2kbp overlapping interval, then computed per-base coverage averaged across each window with bamtocov (Birolo and Telatin 2022). A custom python script was used to pull out genomic coordinates of continuous windows with abnormally high or low coverage depth based on 5^th^ and 95^th^ percentile cutoffs, which we marked as discrepancies for inspection. These regions were marked as discrepancies for further investigation and manual correction where possible. Using this method, we detected and gap-filled a ~500 bp (GAAAA)_n_ tandem repeat region on chr13B by locally assembling the UL reads. Finally, by aligning unplaced contigs against the chromosomal scaffolds, we discovered and manually corrected a ~10 kbp misassembly upstream of the chr13B rDNA array. After each curation step, we aligned duplex reads to the revised assembly to check if alignments agreed with the edits.

**Supplemental Figures S1–S16**

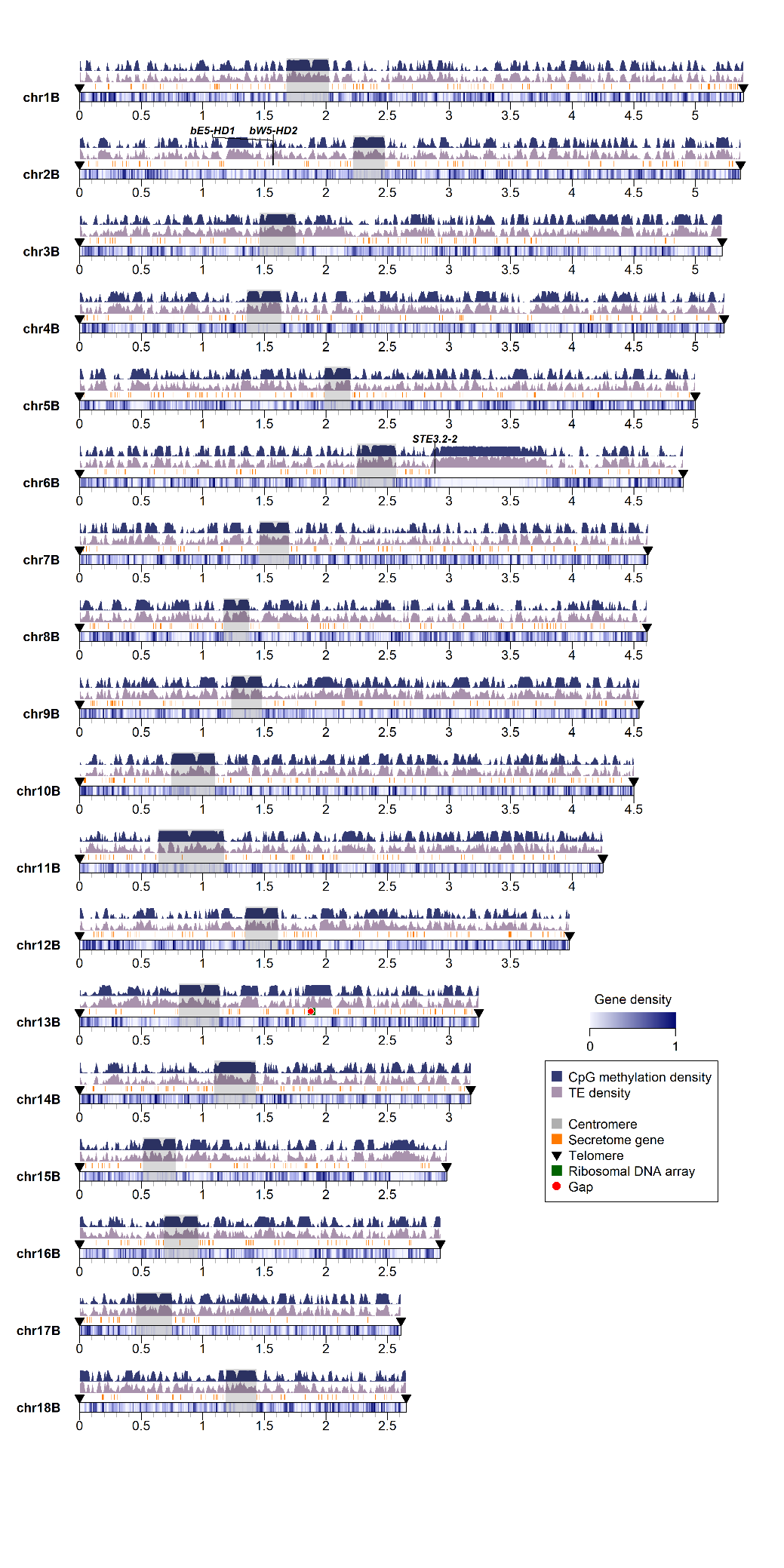

**Supplemental Figure S1.** Karyoplot of the 18 chromosomes of *Pst*104E haplotype B, showing density of CpG methylation and transposable elements (TEs) as peaks, and gene density as heatmaps within chromosome ideograms (10 kbp sliding windows). Locations of centromere, telomeres, secretome genes, ribosomal DNA array and assembly gaps are annotated as per legend.

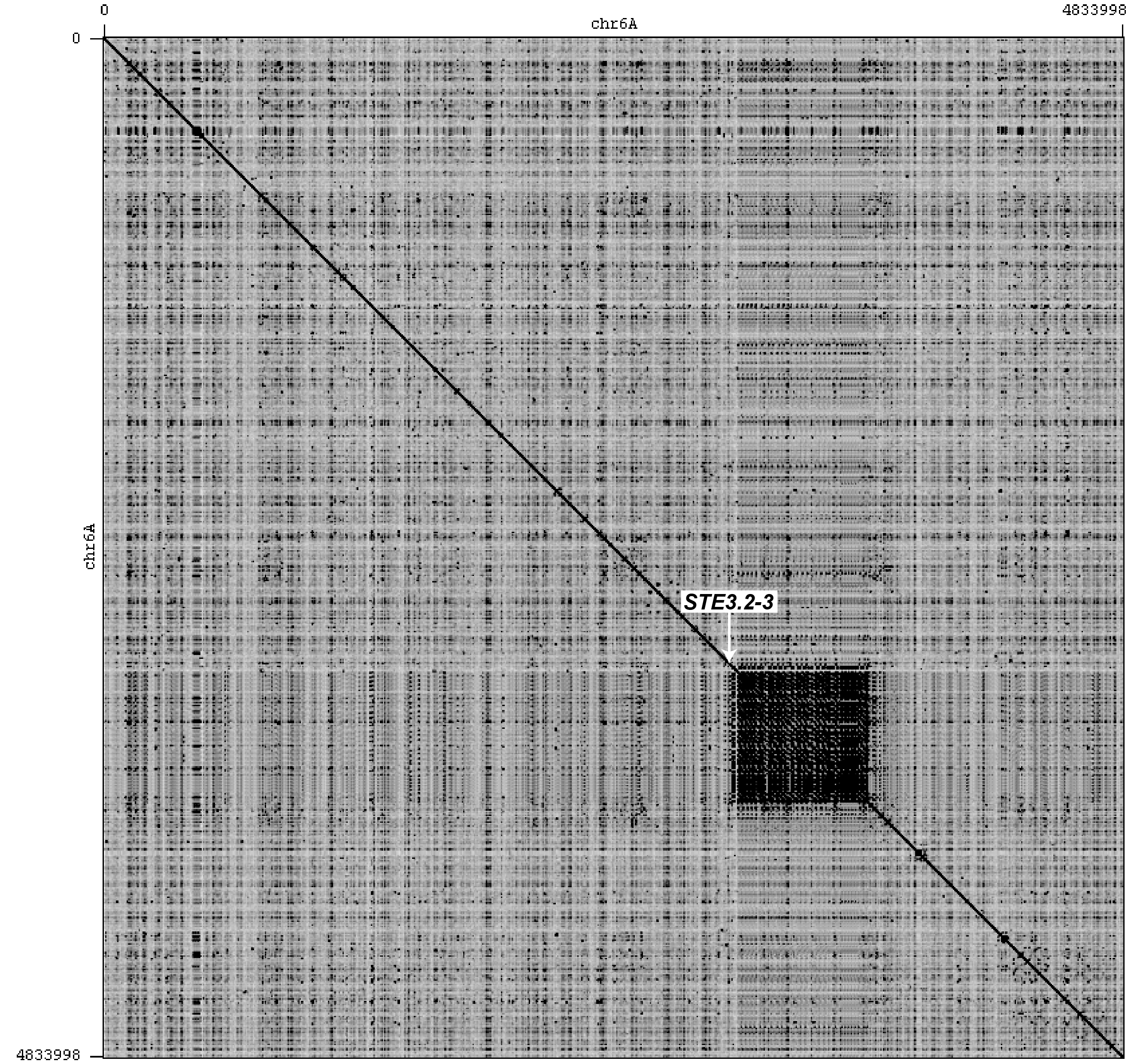

**Supplemental Figure S2.** Self-alignment dotplot of *Pst*104E chromosome 6A. The large dark “patch” spanning ~650 kbp represents the highly repetitive LTR retrotransposon-rich region near the mating type *PR* locus, *STE3.2-3*, as previously documented (Luo et al. 2024). An assembly gap is present in this region potentially due to the high repetitiveness and near-identical repeat units. The corresponding region on haplotype B is completely assembled (not shown). Aligned and visualised in Gepard (Krumsiek et al. 2007).

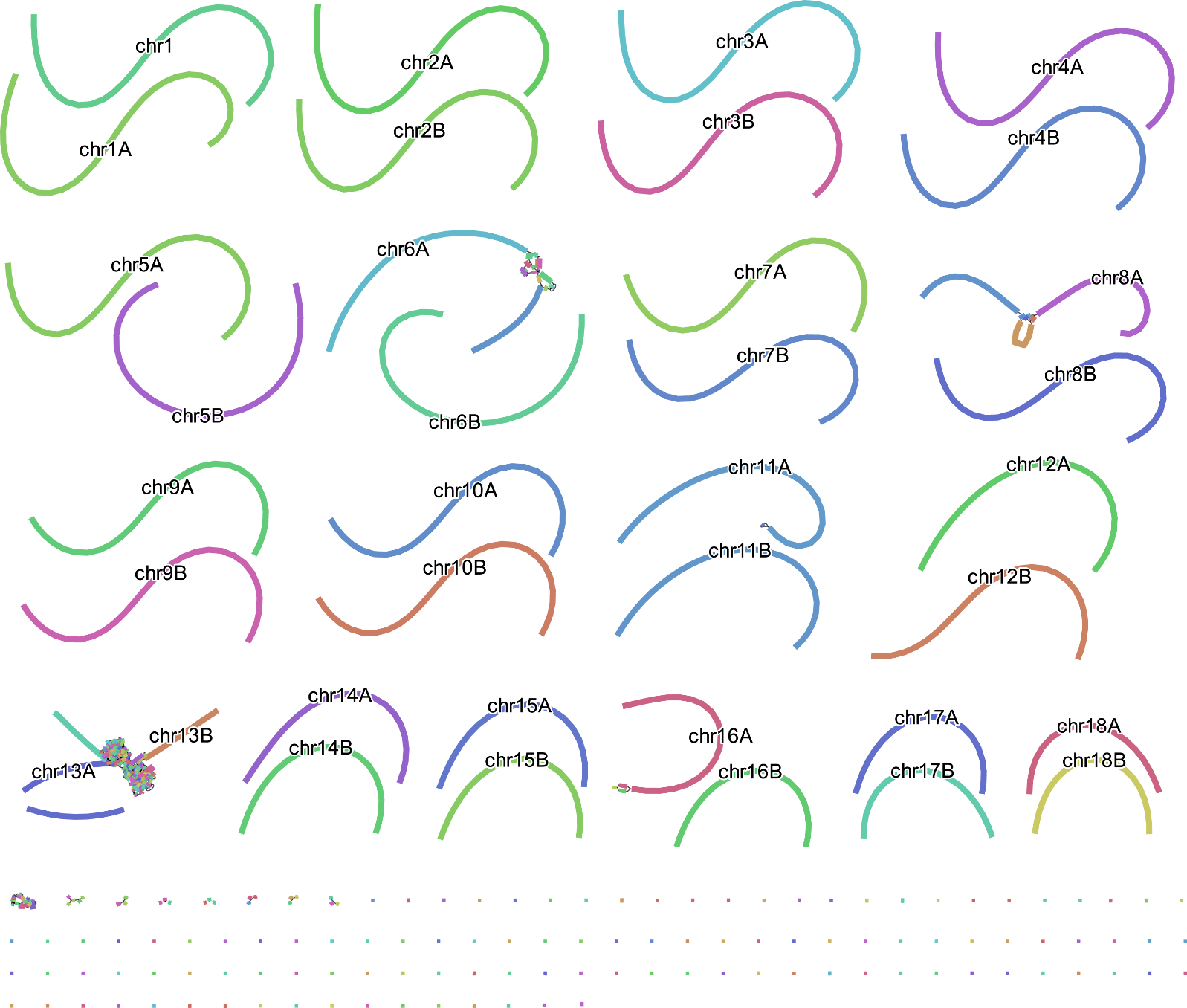

**Supplemental Figure S3.** Genome string graph of raw Verkko assembly of *Pst*104E. Most homologous chromosomes were directly resolved into two linear sequences. Tangles represent ambiguous connections at complex genomic regions, usually associated with long, highly identical repetitive sequences (e.g. rDNA arrays on chromosome 13). All identified assembly gaps coincided with these tangles (one gap on chr6A, chr13A and chr13B; two gaps on chr8A). The small unresolved graph bubble at the chromosomal end of chr16A may indicate a large duplication event at the subtelomeric region leading to an unassembled telomere on its q-arm.

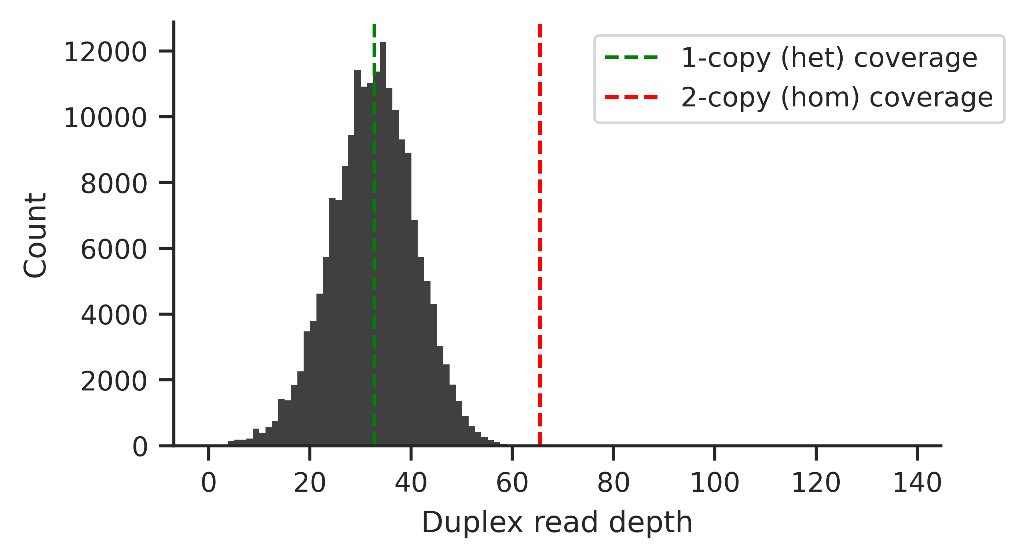

**Supplemental Figure S4.** Histogram of duplex read depth when mapped against the phased *Pst*104E assembly. If an assembly contains haplotype-collapsed regions, the reads derived from both haplotypes would map to the same location giving rise to 2x haploid coverage (red dotted line). Here, a single peak centres at the expected 1x haploid coverage (green dotted line), and no visible peak is present at the expected 2x haploid coverage, implying that the assembly is completely (or almost completely) phased.

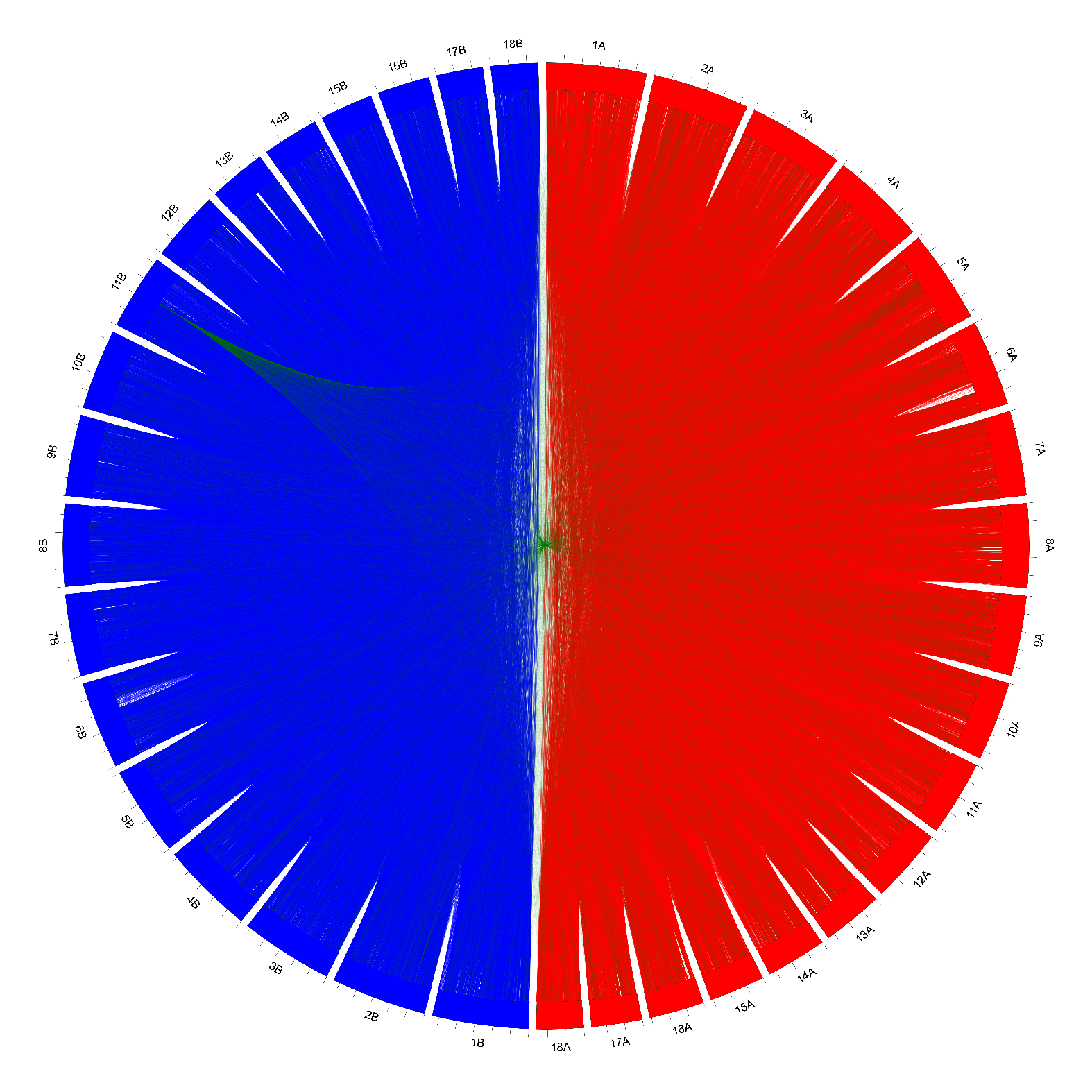

**Supplemental Figure S5.** Circos plot visualising within- and cross-haplotype Hi-C contact links for confirming phasing correctness of the *Pst*104E assembly. Each chromosome sequence was split into 20 kbp genomic windows, for which the rolling means of Hi-C links were computed. Hi-C links mapped within haplotype A are illustrated in red lines; haplotype B in blue lines. Cross-haplotype Hi-C links are illustrated in green lines. The 113^th^ window on chromosome 11B shows frequent cross-haplotype links with haplotype A chromosomes implying a potential minor phase switch left in the assembly.

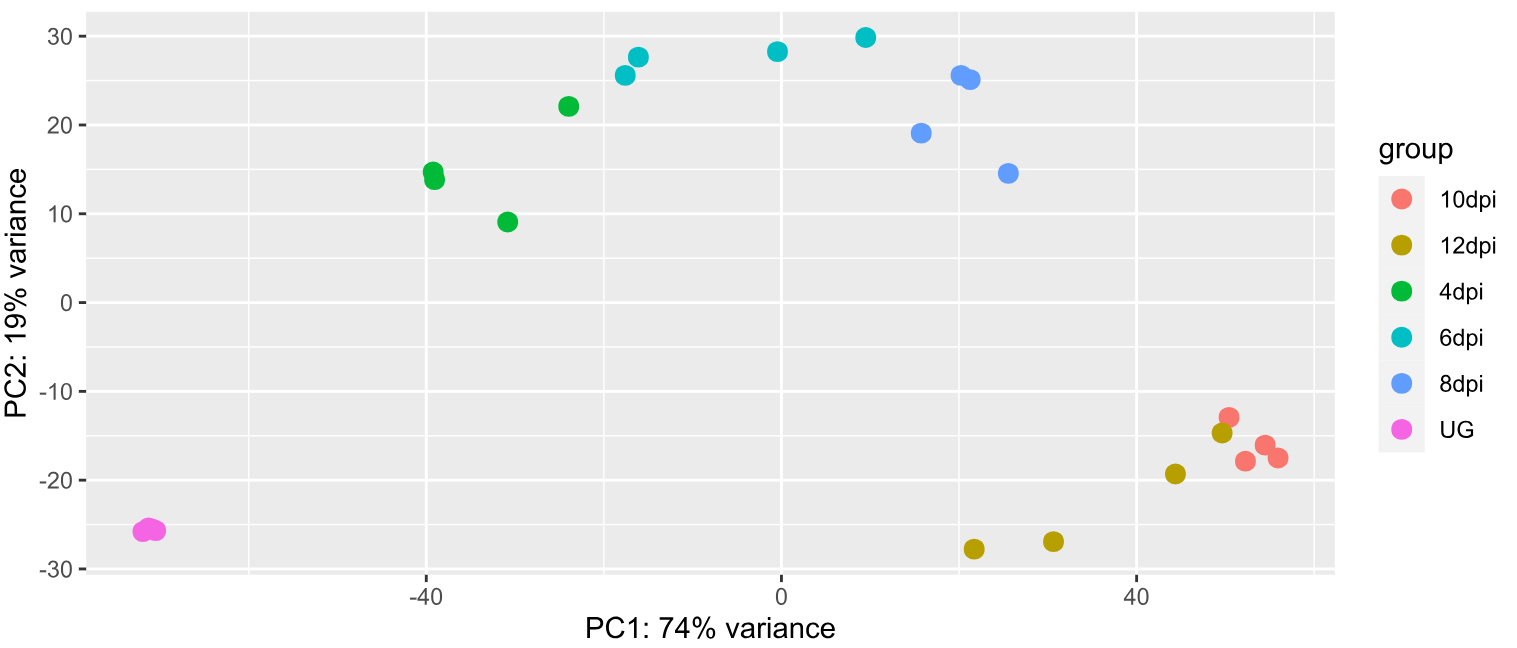

**Supplemental Figure S6.** Principle component analysis plot of the time-course ONT cDNA transcriptome dataset of *Pst*104E. Clear separations could be seen among the transcriptomic profiles derived from ungerminated (dormant) spores, early-mid host infection (4, 6 and 8 dpi) and late-infection (10 and 12 dpi) timepoints. UG: ungerminated urediniospores; dpi: days post infection.

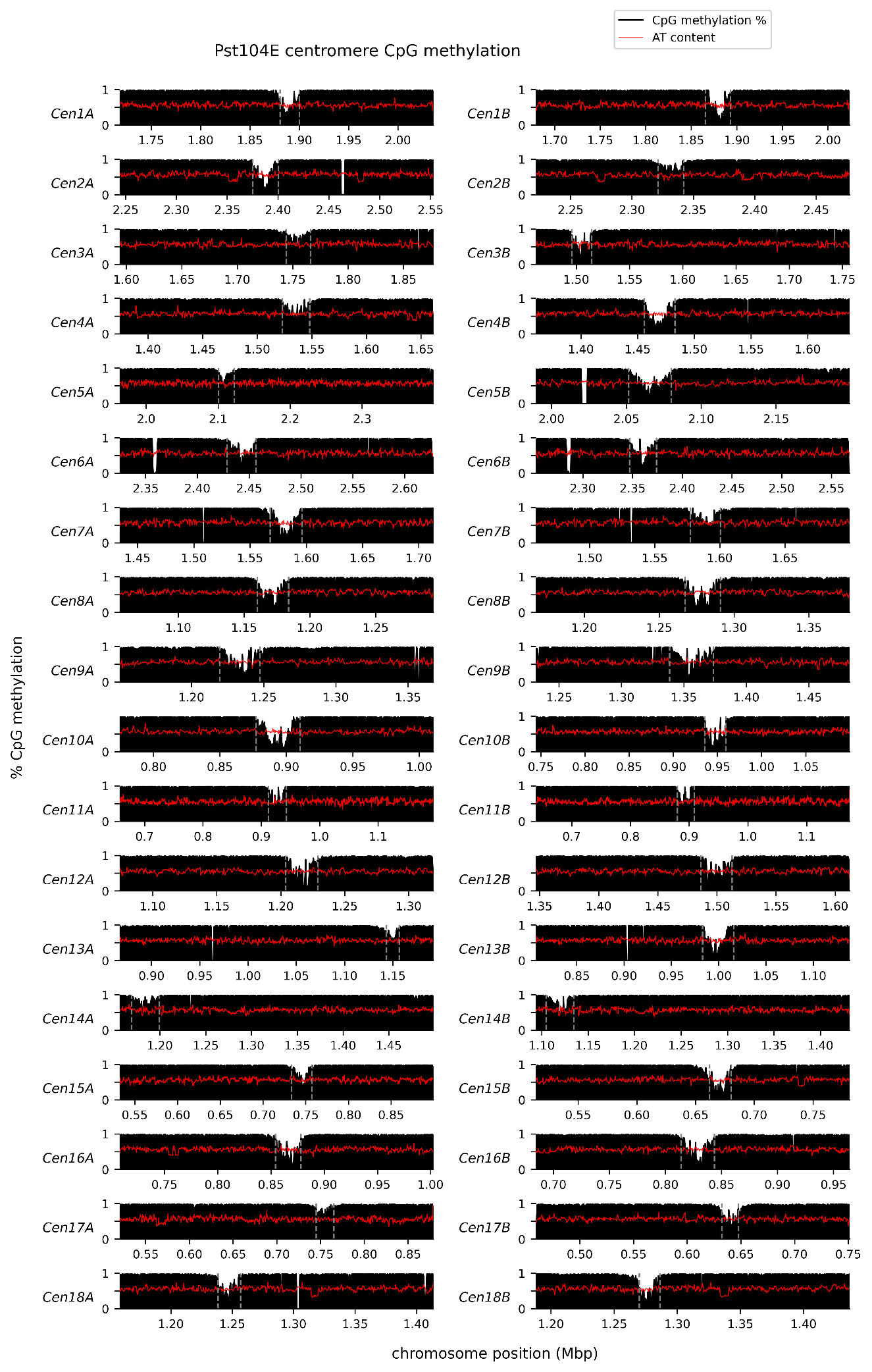

**Supplemental Figure S7.** CpG methylation profiles (black histogram) and % AT content (red line) of all 36 *Pst*104E centromeres. Rolling means were computed over 500 bp windows. A methylation depletion valley with the mean size of ~24.8 Kbp consistently appeared throughout all centromeres, indicating CDR signals (enclosed by grey dotted line).

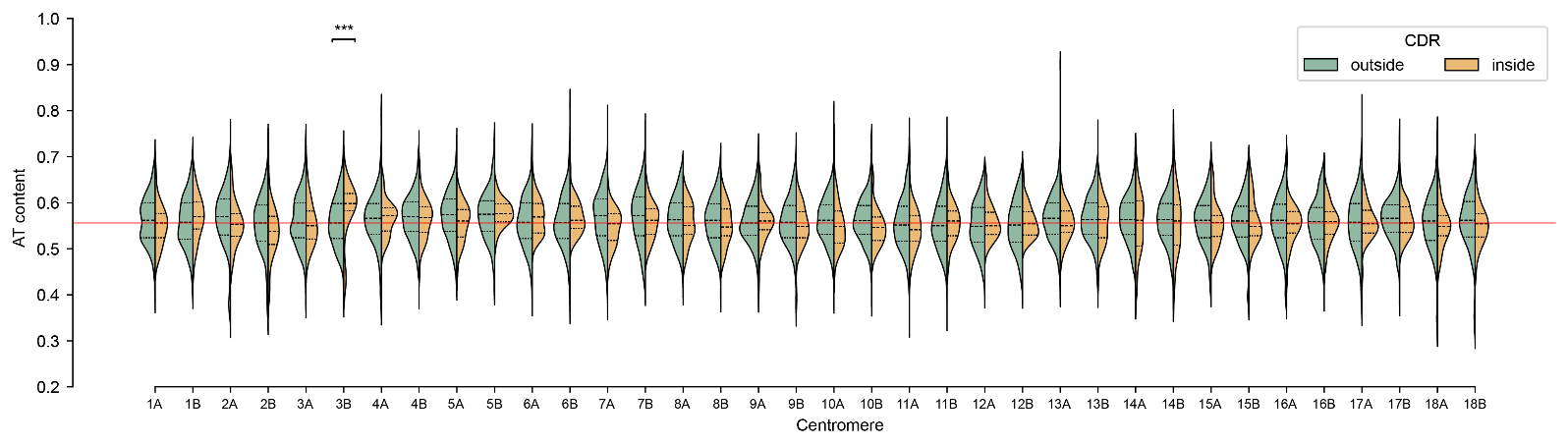

**Supplemental Figure S8.** Split violin plots comparing centromeric AT content in 500 bp windows inside (yellow) and outside (green) of centromere dip region (CDR), which is marked by the CpG methylation depletion signal in otherwise fully methylated centromere. The three dashed lines inside each violin denote the 25^th^, 50^th^ (median) and 75^th^ quartiles. Red solid line plotted at ~0.56 indicates the genome-wide AT content. This was analysed to check whether CDR signals were associated with elevated AT content. One-tailed Mann-Whitney U test (***, p<0.001).

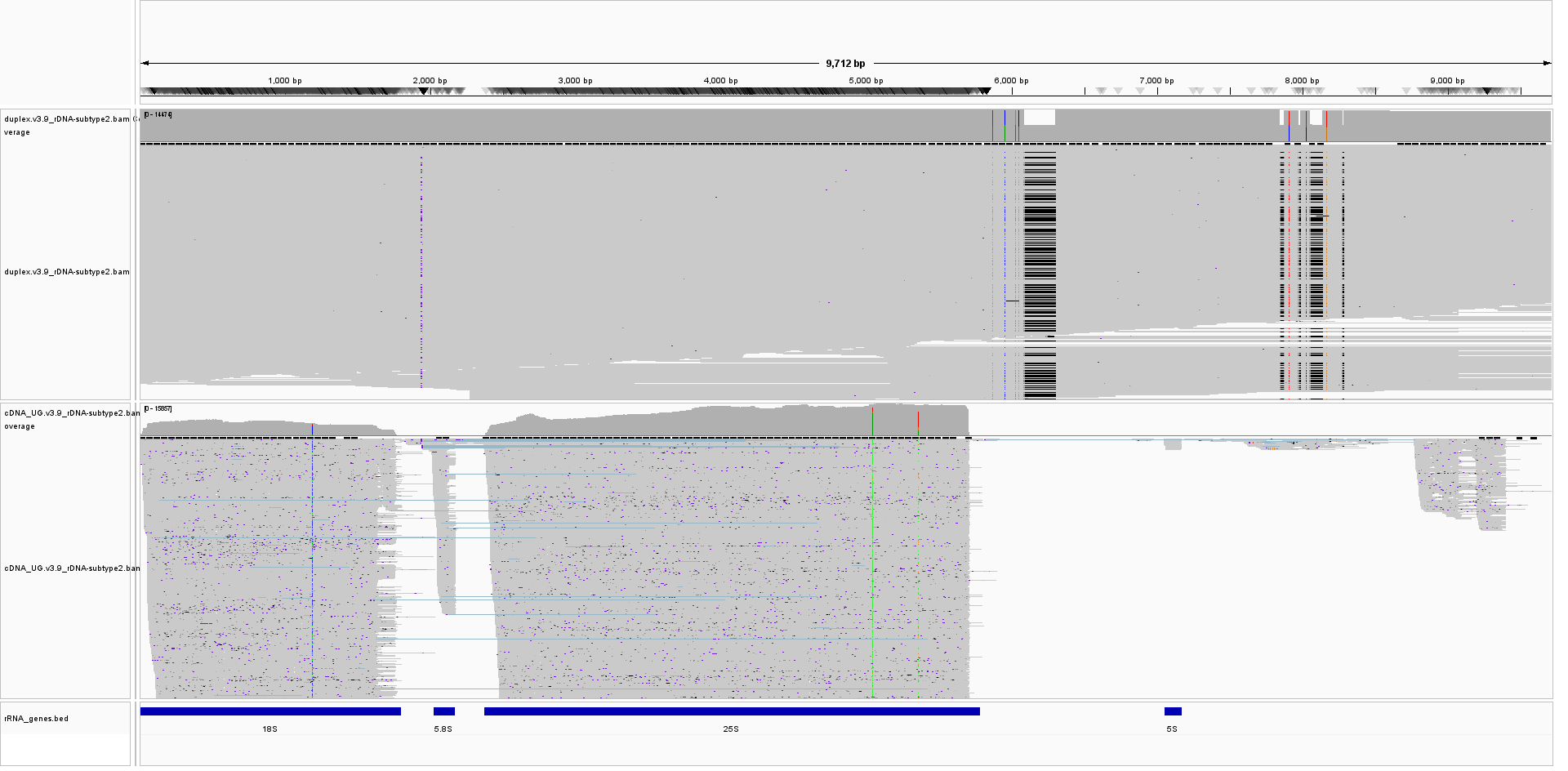
**Supplemental Figure S9.** IGV visualisation of ONT duplex (upper track) and cDNA (lower track) long-read alignments against the canonical rDNA repeat unit of *Pst*104E. Blue tracks at the bottom represent the annotated catalytic rRNA genes (18S, 5.8S, 25S and 5S) which were inferred from cDNA alignment for transcriptional activity, combined with results from homology search against rRNA gene references from a range of *Puccinia* species (see Methods). Duplex read alignment reveals two rDNA subtypes that harbour SNPs and length polymorphisms occurring at near-equal frequencies as reflected in the coverage track. The alternate bases revealed in the cDNA alignment did not line up with any high-confidence SNPs detected in aligned duplex reads. A possible explanation for these “misbasecall” profiles at these bases might be caused by rRNA modifications as described in (Begik et al. 2021).

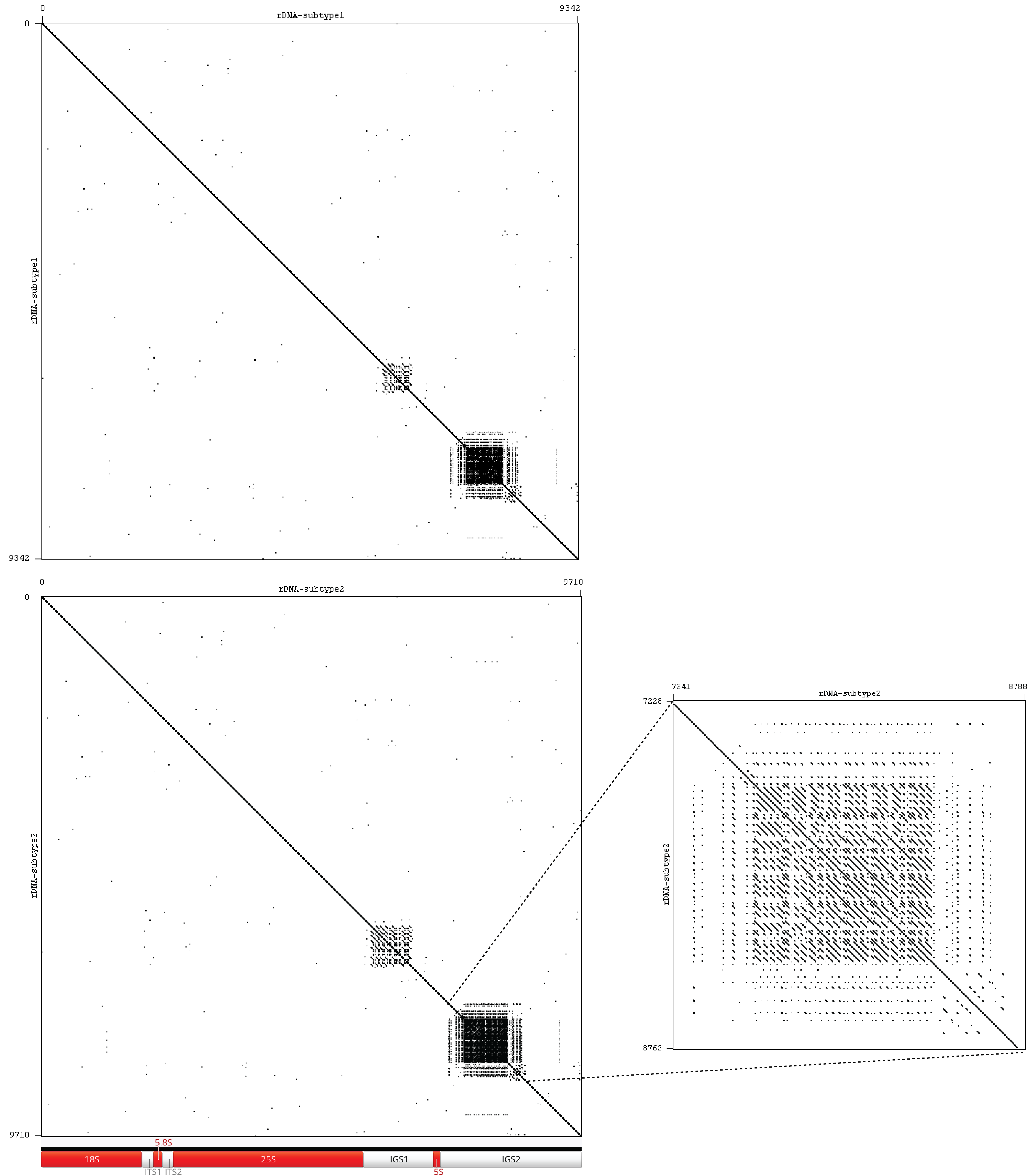

**Supplemental Figure S10.** Self-alignment dotplots of the two major rDNA subtypes (top: subtype #1, bottom: subtype #2) of *Pst*10

4E revealing two subrepeat regions nested within IGS1 and IGS2. Zoomed-in visualisation of the IGS2 subrepeats shows a microsatellite-like structure.

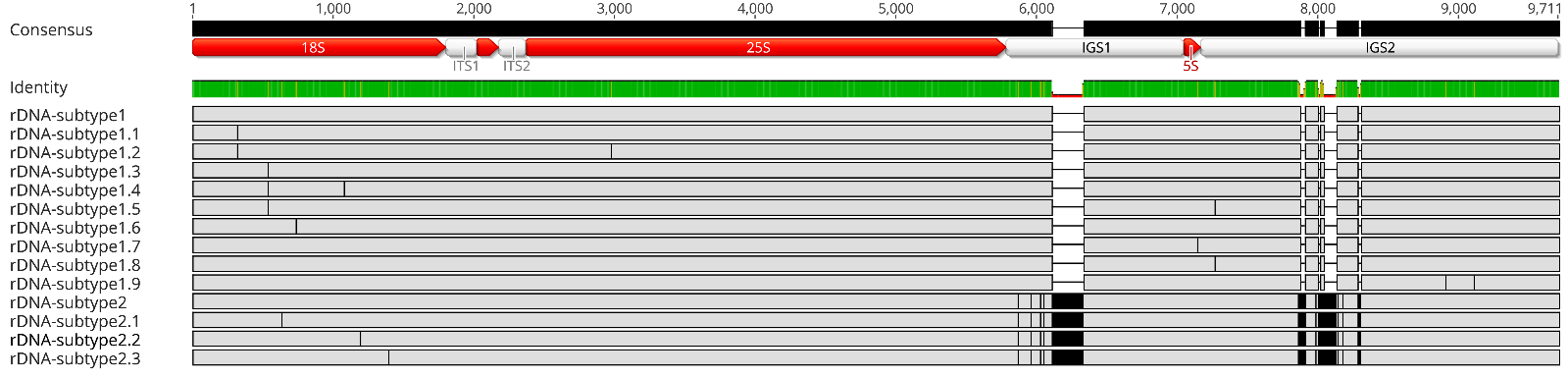

**Supplemental Figure S11.** Multiple sequence alignments of low-frequency variants of the dominant rDNA subtypes #1 and #2, numbered as #1.1–1.9 and #2.1–2.3. High-confidence SNPs were called from duplex read alignments against the canonical rDNA repeat using Bam-readcount (Khanna et al. 2022). SNP combinations were visually determined from aligned reads. Most SNPs accumulated at the 18S gene. Some subtype variants showed overlapping SNPs (e.g. #1.3–1.5) potentially implying spreading point mutations. See Supplemental Table S8 for SNP depths and the inferred copy numbers for each subtype.

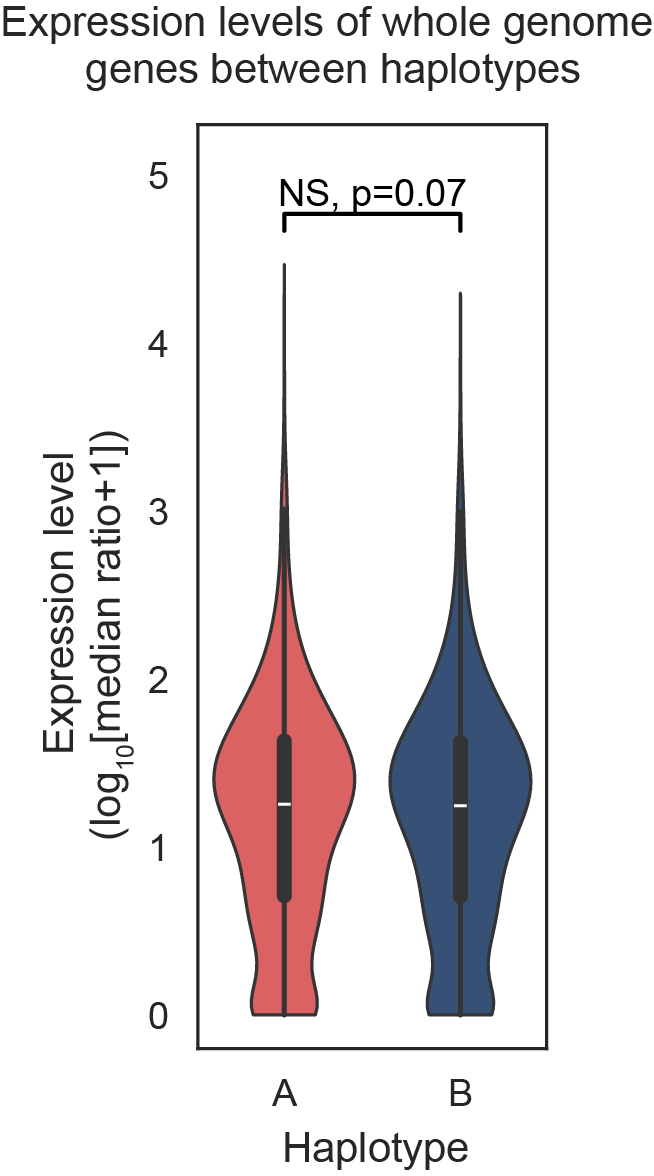

**Supplemental Figure S12.** Violin plots comparing expression level of genome-wide genes (not limited to paired alleles as opposed to Fig. 6B) belonging to the two nuclear haplotype A and B. Mann-Whitney U test (NS, non-significant).

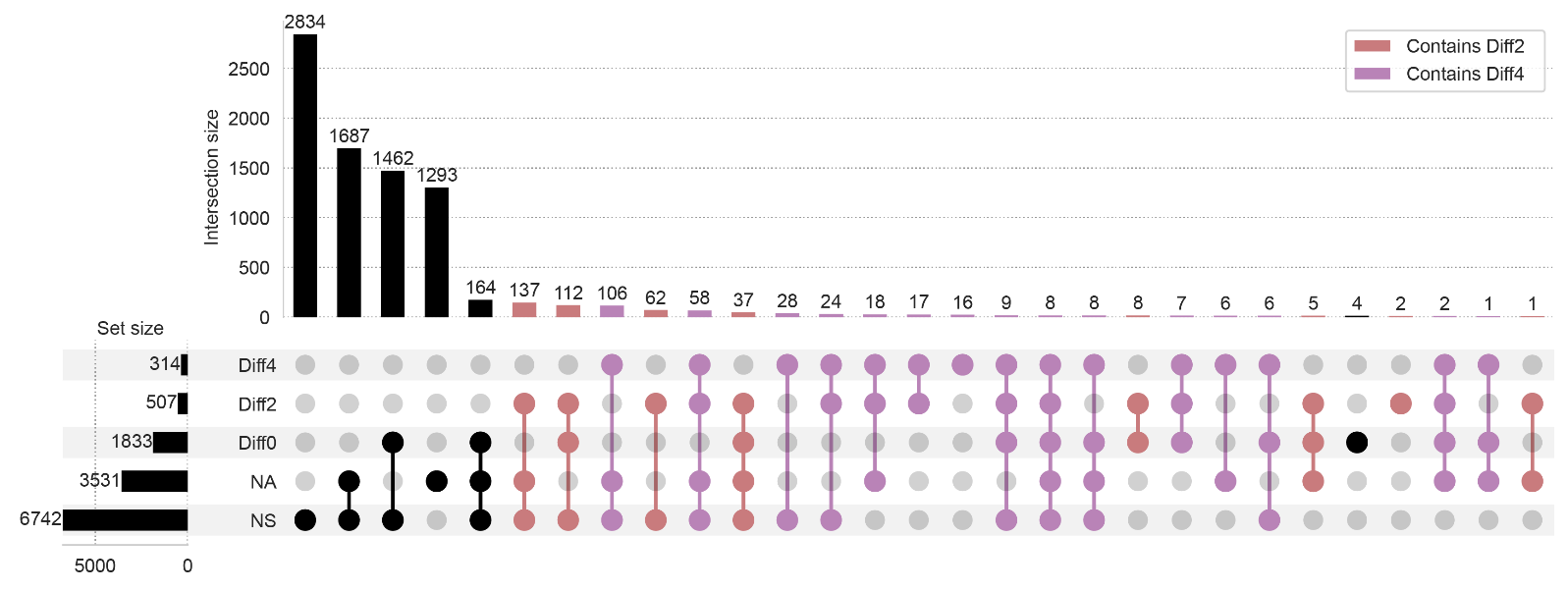

**Supplemental Figure S13.** Upset plot summarising different allele-specific expression (ASE) categories exhibited in all heterozygous biallelic gene pairs (n = 8,122) across the six sampling conditions. ASEs were categorised as followed: (1) no observed expression or no unambiguous transcript mapping at both alleles (NA); (2) no significant difference between alleles with false discovery rate (FDR) adjusted p-value > 0.05 (NS); and (3) significant difference between alleles with adjusted p-value < 0.05, indicating differential ASE. The differential ASE pairs were further classified based on log_2_ fold change (LFC): weak ASE, |LFC|<2 (Diff0); moderate ASE, 2≤|LFC|<4 (Diff2); and strong ASE, |LFC|>4 (Diff4). Highlighted intersections contain allele pairs that exhibited |LFC| greater than two (Diff2 and Diff4) in at least one condition, which were defined as ASE genes for the remainder of the analysis.

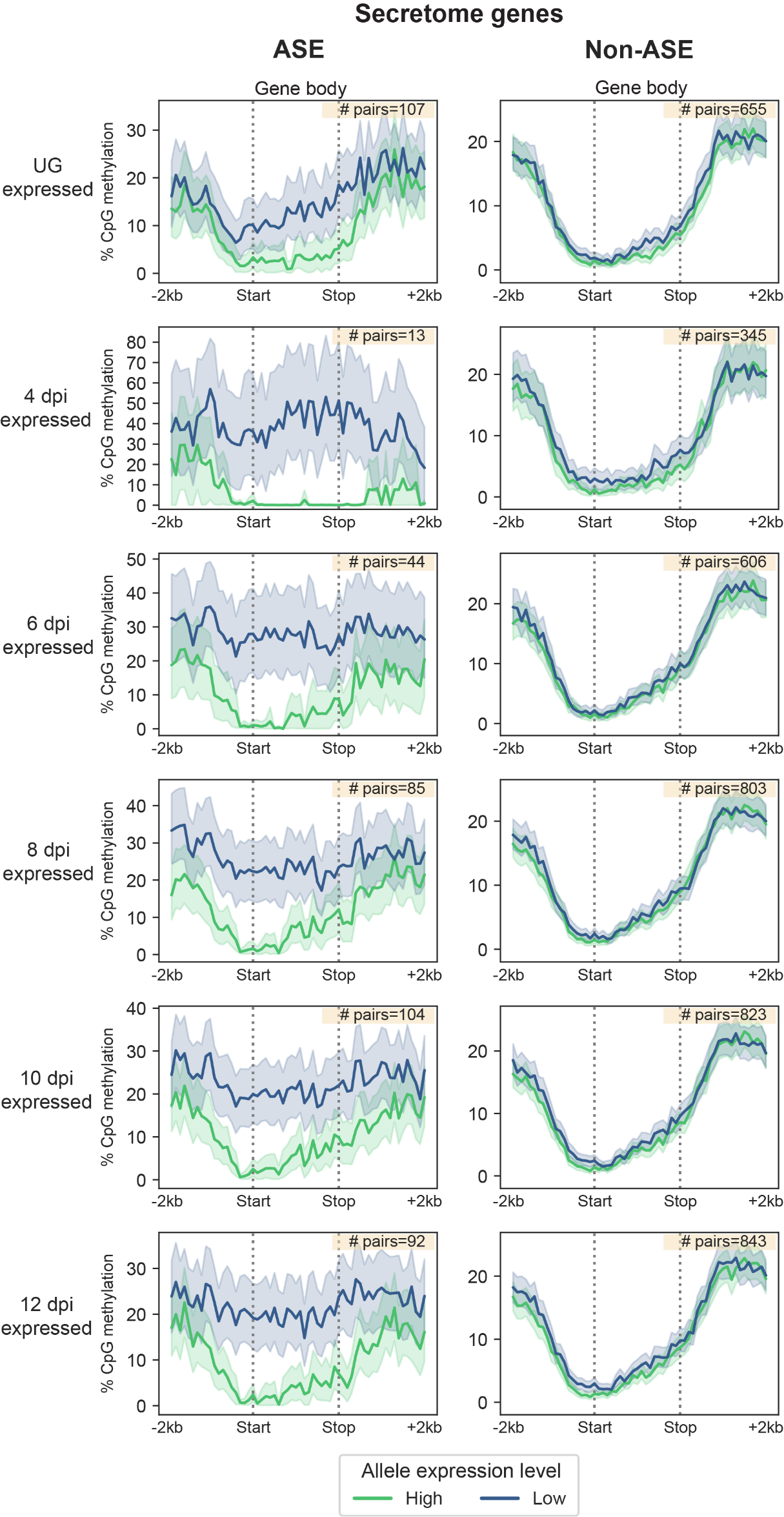

**Supplemental Figure S14.** Distribution of CpG methylation (sampled at UG) across ASE and non-ASE secretome gene bodies including +/- 2 kbp at each of the six sampling conditions. Gene body regions were defined as start and stop codon position, which were extended outwards by 2 kbp to include 5’ upstream and 3’ downstream flanking regions Each region was divided into 20 equally proportioned bins, totalling 60 bins per gene. The per-bin mean percentage coverage CpG methylation is plotted as solid line, with the 95% bootstrapping confidence interval as shaded area. The yellow inset indicates the number of heterozygous allele pairs included.

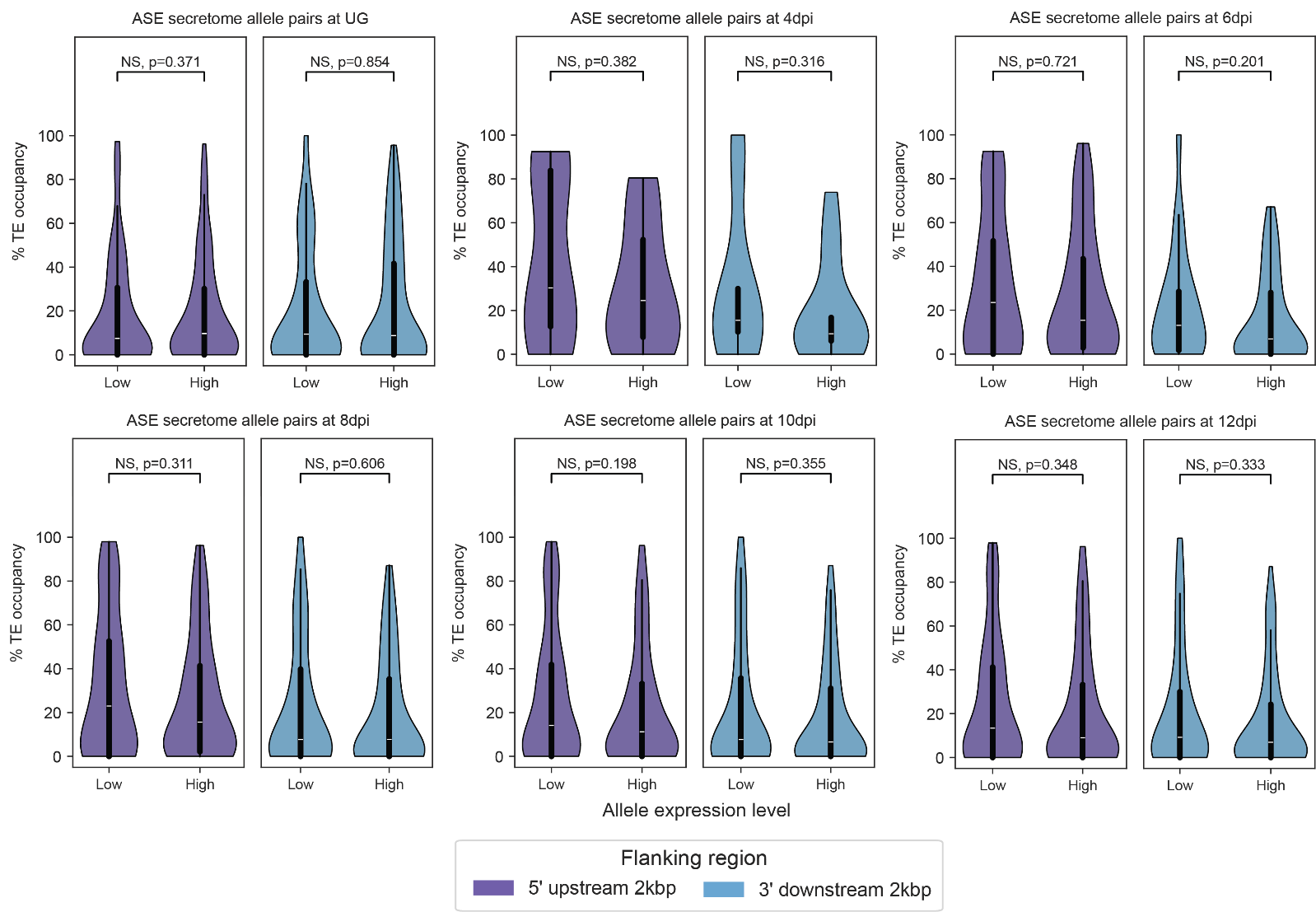

**Supplemental Figure S15.** Violin plots comparing the % TE occupancy between higher- and lower-expressed alleles of ASE secretome genes at their upstream and downstream 2 kbp flanking regions. Mann-Whitney U test (NS, non-significant). No difference in TE occupancy was detected between alleles.

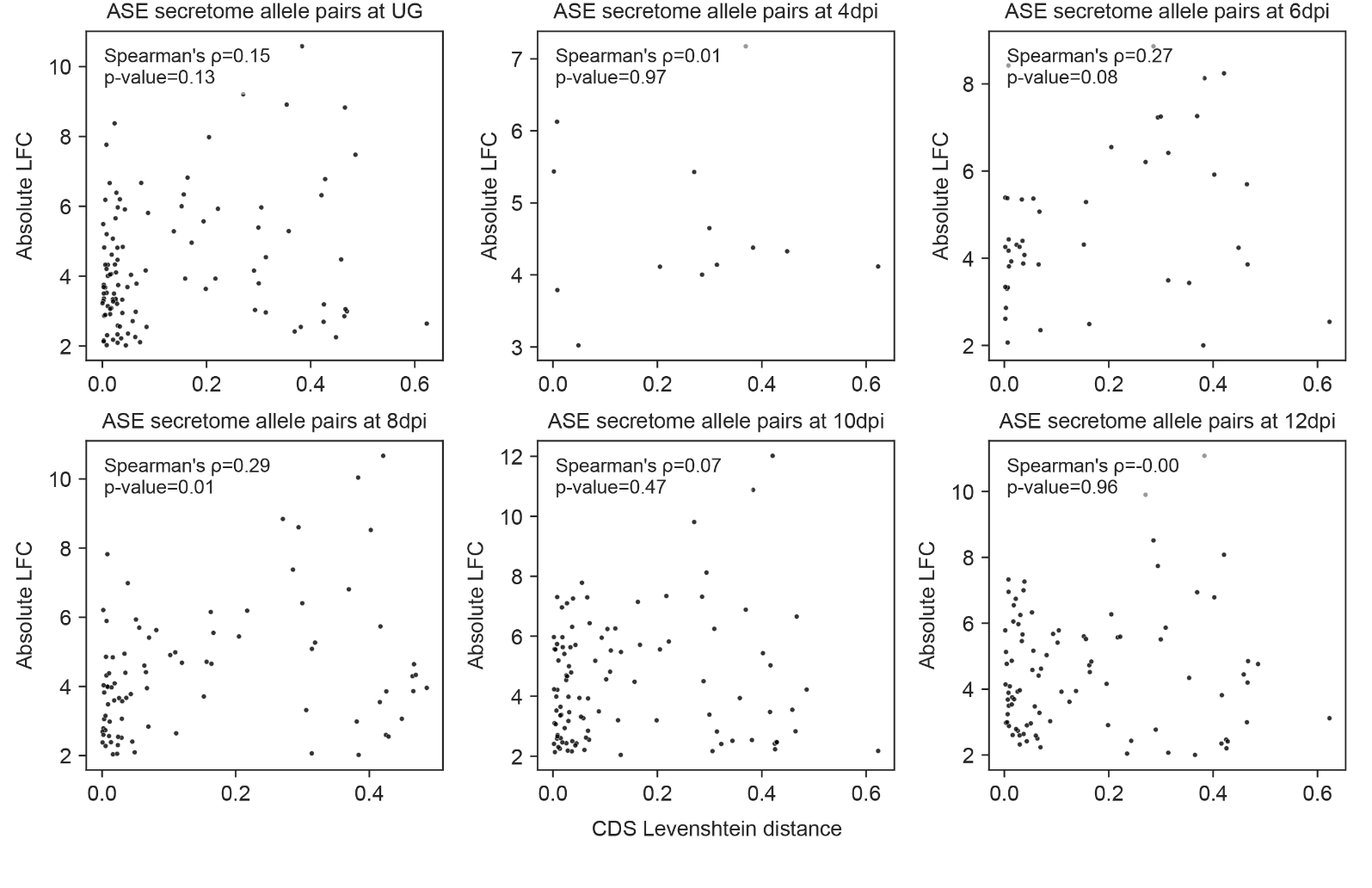

**Supplemental Figure S16.** Spearman’s rho correlation analysis between CDS Levenshtein genetic distance and the expression |LFC| for secretome allele pairs displaying ASE at different conditions. No correlation was detected except for a weak positive correlation shown in the 8 dpi condition.
